# A fluorescent sensor for spatiotemporally resolved endocannabinoid dynamics *in vitro* and *in vivo*

**DOI:** 10.1101/2020.10.08.329169

**Authors:** Ao Dong, Kaikai He, Barna Dudok, Jordan S Farrell, Wuqiang Guan, Daniel J Liput, Henry L Puhl, Ruyi Cai, Jiali Duan, Eddy Albarran, Jun Ding, David M Lovinger, Bo Li, Ivan Soltesz, Yulong Li

## Abstract

Endocannabinoids (eCBs) are retrograde neuromodulators that play an important role in a wide range of physiological processes; however, the release and *in vivo* dynamics of eCBs remain largely unknown, due in part to a lack of suitable probes capable of detecting eCBs with sufficient spatiotemporal resolution. Here, we developed a new eCB sensor called GRAB_eCB2.0_. This genetically encoded sensor consists of the human CB1 cannabinoid receptor fused to circular-permutated EGFP, providing cell membrane trafficking, second-resolution kinetics, high specificity for eCBs, and a robust fluorescence response at physiological eCB concentrations. Using the GRAB_eCB2.0_ sensor, we monitored evoked changes in eCB dynamics in both cultured neurons and acute brain slices. Interestingly, in cultured neurons we also observed spontaneous compartmental eCB transients that spanned a distance of approximately 11 μm, suggesting constrained, localized eCB signaling. Moreover, by expressing GRAB_eCB2.0_ in the mouse brain, we readily observed foot shock-elicited and running-triggered eCB transients in the basolateral amygdala and hippocampus, respectively. Lastly, we used GRAB_eCB2.0_ in a mouse seizure model and observed a spreading wave of eCB release that followed a Ca^2+^ wave through the hippocampus. Thus, GRAB_eCB2.0_ is a robust new probe for measuring the dynamics of eCB release under both physiological and pathological conditions.

---

Cannabis derivatives have long been used for medicinal and recreational purposes across many cultures in formulations such as marijuana and hashish^1^. Bioactive compounds in cannabis, phytocannabinoids, exert their function by “hijacking” the body’s endogenous cannabinoid (endocannabinoid, or eCB) system. The biological function of eCBs—majorly two lipid metabolites 2-arachidonoylglycerol (2-AG) and anandamide (AEA)—is primarily mediated by the activation of type1 and type 2 cannabinoid receptors (CB1R and CB2R)^2^. eCBs are widely distributed throughout the peripheral and central nervous system, where they serve as important neuromodulators. Interestingly, unlike other classical neurotransmitters stored in synaptic vesicles and released from the presynaptic terminal, eCBs are typically produced and released from the postsynaptic compartment in a neuronal activity-dependent manner, then retrogradely travel to the presynaptic terminal and activate the CB1R, activation of which often results in an inhibition of presynaptic neurotransmitter release^3,4^. In addition, eCBs also play a role in glial cells and in intracellular organelles^5–9^. In the brain, eCBs participate in the short-term and long-term synaptic plasticity of glutamatergic and gamma-aminobutyric acid (GABA)-ergic synapses in a variety of regions, including the cerebral cortex, hippocampus, striatum, ventral tegmental area, amygdala and cerebellum^4,10^, playing an important role in a wide range of physiological processes such as development, emotional state, pain, the sleep/wake cycle, energy metabolism, reward, and learning and memory^11–15^. Given the broad distribution and variety of functions of eCBs, dysregulation of the eCB system has been associated with a plethora of disorders, including neuropsychiatric and neurodegenerative diseases, epilepsy, cancer, and others^16–18^. The eCB system has therefore emerged as a promising target for treating neurological diseases^19,20^.

Although we know much about the eCB biochemistry and physiology, the spatiotemporal dynamics of eCB release in the brain remains largely unknown. Synaptic transmission mediated by classic neurotransmitters such as glutamate and GABA and their respective ionotropic receptors can occur in a timescale on the order of milliseconds and is generally spatially confined to the synaptic cleft in the nanometer range^21^. In contrast, signaling via endocannabinoid receptors is believed to last on the order of seconds and over a distance on the order of tens of microns. However, this assumption has not been tested directly, largely because existing methods for measuring eCB signaling lack the necessary spatiotemporal resolution. For example, although qualitative and quantitative measurement of eCBs in brain tissues can provide valuable information regarding eCB levels, these measurements usually require the extraction, purification and analysis of lipids by chromatography and mass spectrometry^22,23^, therefore, this approach has poor spatial and temporal resolution and cannot be used to measure eCBs *in vivo*. Another approach is electrophysiology coupled with pharmacology and/or genetics, which is often used to indirectly measure eCB activity by measuring eCB-mediated synaptic modulation^24–27^; however, this method is mostly used in *in vitro* preparations and has relative low spatial resolution. Another method microdialysis, while challenging for hydrophobic lipid molecules, has been used to monitor eCB abundance in the brain during pharmacological manipulations and behaviors^28,29^, but it has a long sampling interval (at least 5 minutes) that is well beyond the time scale of synaptic plasticity mediated by eCBs (~sub-second to seconds), preventing the accurate detection of eCBs in real time *in vivo*. Therefore, development of an *in vivo* eCB detection tool with satisfactory spatiotemporal resolution would meet a clear need in this field.

Recently, our group and others developed a series of genetically-encoded tools for sensing neurotransmitters and neuromodulators based on G protein-coupled receptors (GPCRs) and circular-permutated (cp) fluorescent proteins^30–38^. Using this highly successful strategy, we developed a novel GPCR activation-based (GRAB) eCB sensor called GRAB_eCB2.0_ (or simply eCB2.0) based on the human CB1R and cpEGFP. The eCB2.0 sensor has high specificity for eCBs, kinetics on the order seconds, and a fluorescence response of approximately 800% to 2-AG and 550% to AEA, respectively. After validating the *in vitro* performance of eCB2.0 in both cultured cells and acute brain slices, we then expressed the sensor in mice and reliably monitored foot-shock evoked eCB signals in the basolateral amygdala in freely moving mice and eCB dynamics in the mouse hippocampus during running and seizure activity.

## RESULTS

### Development and *in vitro* characterization of GRAB_eCB_ sensors

Among the two eCB receptors, we chose CB1R as the scaffold for developing a GRAB eCB sensor, as this receptor has a higher affinity for eCBs than CB2R^39^. We first inserted the intracellular loop 3 (ICL3)-cpEGFP module of our recently developed GRAB_NE_ sensor^33^ into the corresponding ICL3 in the human CB1R (**Fig. 1a**). After screening various insertion sites and GRAB_NE_ ICL3 truncation constructs, we generated the first-generation eCB sensor called GRAB_eCB1.0_ (eCB1.0), which showed a moderate response (100% increase in fluorescence) to ligand and an apparent affinity of 3 μM for 2-AG (**Fig. 1b and Extended Data Fig. 1a**). To improve the dynamic range of our eCB sensor, we then selected 8 residues in cpEGFP for individual randomized mutation based our the experience gained through the development of previous GRAB sensors^30,32–34,36–38^ (**Extended Data Fig. 1b**). Combining several single-mutation candidates—each with improved performance—resulted in the GRAB_eCB1.5_ sensor (eCB1.5), which has a 2-fold higher response than eCB1.0 (**Extended Data Fig. 1a**). We next focused on the receptor’s ligand binding pocket in order to further improve the sensor’s dynamic range and affinity. The residues F177^2.64^, V196^3.32^ and S38 3^7.39^ were selected for targeted screening based on the studies of CB1R structure^40–45^ (**Extended Data Fig. 1c**). Interestingly, we found that introducing the S383^7.39^T mutation in eCB1.5 produced an increased response to 2-AG with a similar apparent affinity, whereas adding the F177^2.64^A mutation abolished the response to 2-AG (**Extended Data Fig. 1a**). We therefore selected the eCB1.5 with the S383^7.39^T mutation as the second-generation GRAB_eCB2.0_ sensor (eCB2.0), and eCB1.5 with both the S383^7.39^T and F177^2.64^A mutations as a non-responsive negative control, which we call GRAB_eCBmut_ sensor (eCBmut) (**Extended Data Fig. 2**).

**Fig. 1.**
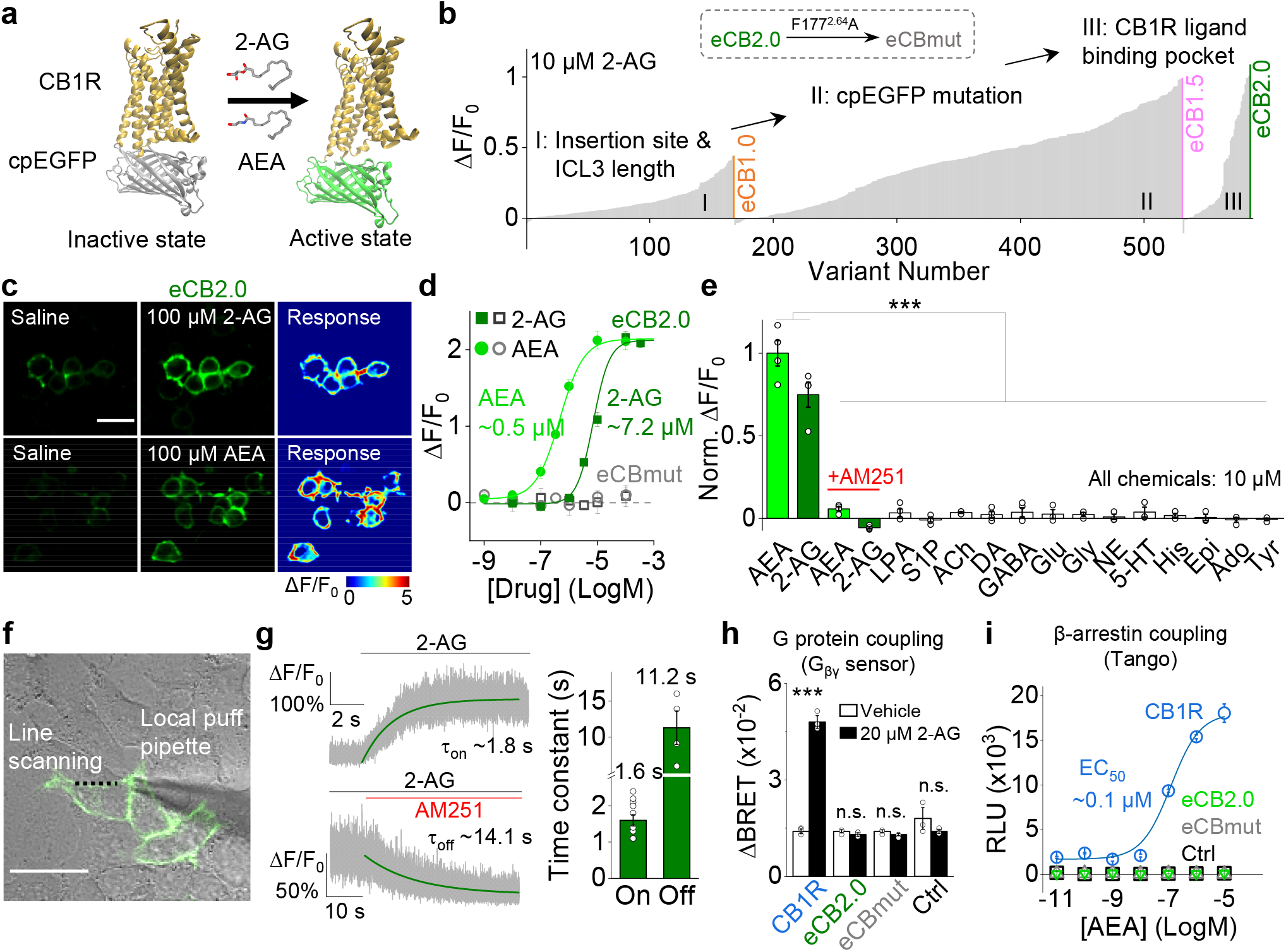
Development, optimization, and characterization of GRAB_eCB_ sensors in HEK293T cells. **a**, Schematic diagram depicting the design and principle of the GRAB_eCB_ sensor, consisting of the CB1 receptor and circular-permutated GFP. Ligand binding activates the sensor, inducing a change in fluorescence. **b**, Screening and optimization steps of GRAB_eCB_ sensors and the normalized fluorescence response to 10 μM 2-AG. eCBmut was generated by introducing the F177^2.64^A mutation in eCB2.0. **c**, Expression and fluorescence change in response to 100 μM 2-AG and AEA in HEK293T cells expressing eCB2.0. Scale bar, 30 μm. **d**, Dose-response curves measured in HEK293T cells expressing eCB2.0 or eCBmut, with the corresponding EC_50_ values for 2-AG and AEA shown; n = 3 wells each. **e**, Normalized fluorescence change in response to the indicated compounds (each at 10 μM) measured in cells expressing eCB2.0; n = 3–4 well each. Where indicated, the CB1R inverse agonist AM251 was also added. LPA, lysophosphatidic acid; S1P, sphingosine-1-phosphate; ACh, acetylcholine; DA, dopamine; GABA, gamma-aminobutyric acid; Glu, glutamate; Gly, glycine; NE, norepinephrine; 5-HT, 5-hydroxytryptamine; His, histamine; Epi, epinephrine; Ado, adenosine; Tyr, tyramine. **f**, Illustration of the localized puffing system using a glass pipette containing 100 μM 2-AG and/or AM251 positioned above an eCB2.0-expressing cell. The dotted black line indicates the region of interest for line scanning. Scale bar, 30 μm. **g**, Change in eCB2.0 fluorescence was measured in an eCB2.0-expressing cell using line scanning; where indicated, 2-AG and AM251 were puffed on the cell. The graph at the right summarizes the on and off time constants measured upon application of 2-AG and upon application of AM251, respectively; n = 11 (t_on_) and 4 (τoff) cells. **h**, G protein coupling was measured using a BRET G_βγ_ sensor in cells expressing CB1R, eCB2.0, or eCBmut. **i**, β-arrestin coupling was measured using the Tango assay in cells expressing CB1R, eCB2.0, or eCBmut. Student’s *t* tests were performed in **e** and **h**: ***p < 0.001; n.s., not significant.

When expressed in HEK293T cells, both the eCB2.0 and eCBmut sensors trafficked to the cell membrane (**Fig. 1c**). Upon ligand application, eCB2.0 had a concentration-dependent fluorescence increases to both 2-AG and AEA, with a maximum response of approximately 2 fold relative to baseline and the half maximal effective concentrations (EC_50_) for 2-AG and AEA of 7.2 μM and 0.5 μM, respectively; in contrast, eCBmut showed no response to 2-AG or AEA at all concentrations tested (**Fig. 1d**). We then tested whether the sensor’s response is specific to eCBs compared to other neurotransmitters. We found that eCB2.0 responded robustly to both 10 μM AEA and 2-AG, and the response was abolished by the CB1R inverse agonist AM251; moreover, no other neurotransmitters or neuromodulators tested elicited a response in cells expressing eCB2.0 (**Fig. 1e**).

Next, we measured the kinetics of eCB2.0 signaling using a rapid localized solution application system in which compounds were puffed directly on the cell (**Fig. 1f**). To measure the onset rate (t_on_), 100 μM 2-AG was puffed on eCB2.0 expressing cell; to measure the offset rate (t_off_), 100 μM AM251 was puffed in the presence of 10 μM 2-AG. Using this approach, we measured averaged t_on_ and t_off_ values of 1.6 s and 11.2 s, respectively (**Fig. 1g**). To examine whether eCB sensors couple with intracellular signaling pathways, we measured G-protein activation using a G_βγ_ bioluminescence resonance energy transfer (BRET) sensor based on the G_βγ_ binding region of phosducin fused to NanoLuc luciferase. This unified BRET sensor was based upon similar systems^46,47^. Treating cells expressing CB1R with 2-AG induced a robust increase in BRET, consistent with G protein activation; in contrast, 2-AG had no effect on BRET in mock-transfected control cells or in cells expressing either eCB2.0 or eCBmut (**Fig. 1h**). We also measured Δ-arrestin recruitment using the Tango GPCR assay^48^ and found that AEA induced a robust, concentration-dependent response in cells expressing CB1R but had no effect in control cells or cells expressing either eCB2.0 or eCBmut (**Fig. 1i**). Taken together, these data indicate that our eCB2.0 sensor binds eCBs but does not couple to downstream effector proteins and therefore likely does not affect cellular physiology.

We then examined the expression pattern of the eCB sensor in neurons by sparsely expressing eCB2.0 in cultured rat cortical neurons. We found that eCB2.0 trafficked to the entire neuronal cell membrane, including the axons and dendrites, as shown by colocalization with the axonal presynaptic marker synaptophysin-mScarlet and the postsynaptic marker PSD95-mScarlet (**Fig. 2a**). To measure the response of eCB2.0 in neurons, we infected cultured rat cortical neurons using an adeno-associated virus (AAV) expressing either eCB2.0 or eCBmut under the control of the human *SYN1* (synapsin) promoter to drive expression in all neurons (**Fig. 2b**). We found that both 2-AG and AEA elicited concentration-dependent fluorescence responses in neurons expressing eCB2.0, with a maximum fluorescence increase of 800% and 550%, respectively, and an EC_50_ value of 17.2 μM and 0.7 μM, respectively; in contrast, neither 2-AG nor AEA elicited a response in neurons expressing eCBmut, even at 100 μM (**Fig. 2b,c**). We also found that eCB2.0 responses in neurites were higher than in somata (**Fig. 2d**). Finally, bath application of the CB1R agonist WIN55212-2—which can activate eCB2.0 in HEK293T cells (**Extended Data Fig. 3a**)—to eCB2.0-expressing neurons induced a fluorescence increase that was stable for up to 2 hours and blocked completely by AM251 (**Fig. 2e**), suggesting that the sensor does not undergo arrestin-mediated internalization or desensitization and can be used for long-term monitoring of eCB activity.

**Fig. 2.**
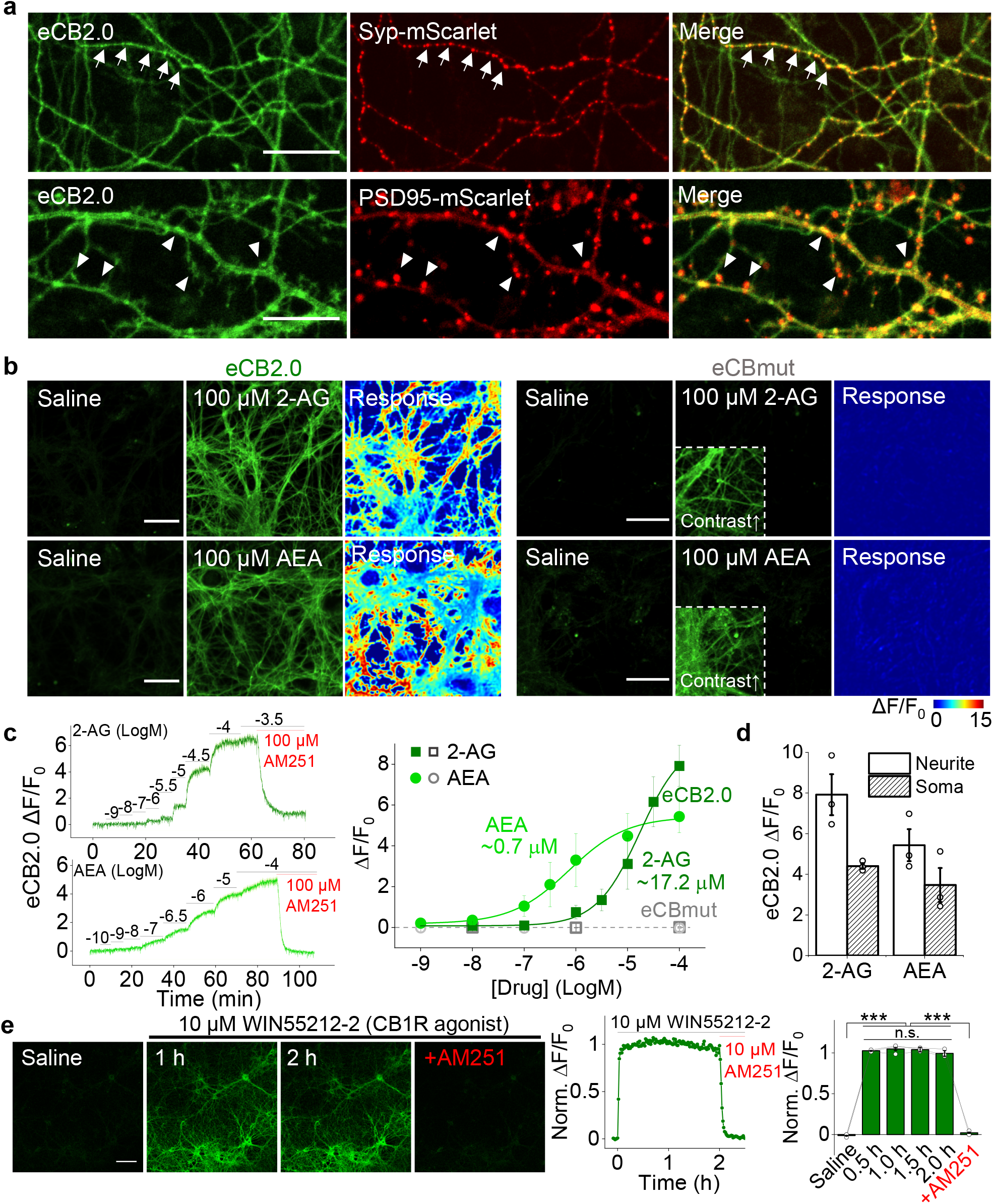
Characterization of GRAB_eCB_ sensors in primary cultured neurons. **a**, Fluorescence microscopy images of primary cultured rat cortical neurons expressing eCB2.0 (green) and either synaptophysin-mScarlet (top row; red) or PSD95-mScarlet (bottom row; red). In the top row, arrows indicate axons; in the bottom row, arrowheads indicate dendrites and dendritic spines. Scale bars, 30 μm (top row) and 15 μm (bottom row). **b**, Fluorescence microscopy images and fluorescence response to 100 μM 2-AG (top row) or AEA (bottom row) in neurons expressing eCB2.0 (left) or eCBmut (right). The insets in the eCBmut images are contrast-enhanced to show expression of the sensor. Scale bars, 30 μm. **c**, (Left) example traces of ΔF/F_0_ measured in an eCB2.0-expressing neuron; the indicated concentrations of 2-AG and AEA, followed by 100 μM AM251, were applied. (Right) dose-response curves measured in neurons expressing eCB2.0 or eCBmut, with the corresponding EC_50_ values shown; n = 3 cultures each. **d**, Summary of the change in eCB2.0 fluorescence in response to 100 μM 2-AG or AEA measured in the neurites and soma; n = 3 cultures each. **e**, Example images (left), trace (middle), and quantification (right) of the change in eCB2.0 fluorescence in response to a 2-hour application of WIN55212-2, followed by AM251; n = 3 cultures each. Scale bar, 100 μm. Student’s *t* test and one-way ANOVA were performed in **e**: ***p < 0.001; n.s., not significant.

### eCB2.0 can be used to measure endogenous eCBs in primary cultured neurons

Cultured neurons are commonly used for studying eCB mediated synaptic modulation^27,49^. We therefore examined whether our eCB2.0 sensor can be used to detect the release of endogenous eCB in cultured rat cortical neurons expressing eCB2.0 together with a red glutamate sensor R^ncp^-iGluSnFR^50^. Applying electrical field stimuli (100 pulses at 50 Hz) elicited robust eCB and glutamate signals (**Fig. 3a**), demonstrating that eCB2.0 can reliably report endogenous eCB release and is compatible with red fluorescent indicators. We then expressed eCB2.0 in neurons loaded with a red fluorescent Ca^2+^ dye Calbryte-590 in order to simultaneously measure eCB release and changes in intracellular Ca^2+^. Applying 100 field stimuli at 50 Hz elicited robust responses with respect to both intracellular Ca^2+^ and eCB release (**Fig. 3b**). Moreover, the rise and decay kinetics of the calcium signal were faster than those of the eCB signal, consistent with the notion that eCB release requires neuronal activity^51^. We also found a strong correlation between the peak Ca^2+^ signal and the peak eCB signal when applying increasing numbers of stimuli (R^2^ = 0.99, **Fig. 3c**); importantly, in the absence of extracellular Ca^2+^, even 20 pulses were unable to elicit either a Ca^2+^ signal or an eCB2.0 response (**Fig. 3c**), confirming the requirement of calcium activity on eCB release.

**Fig. 3.**
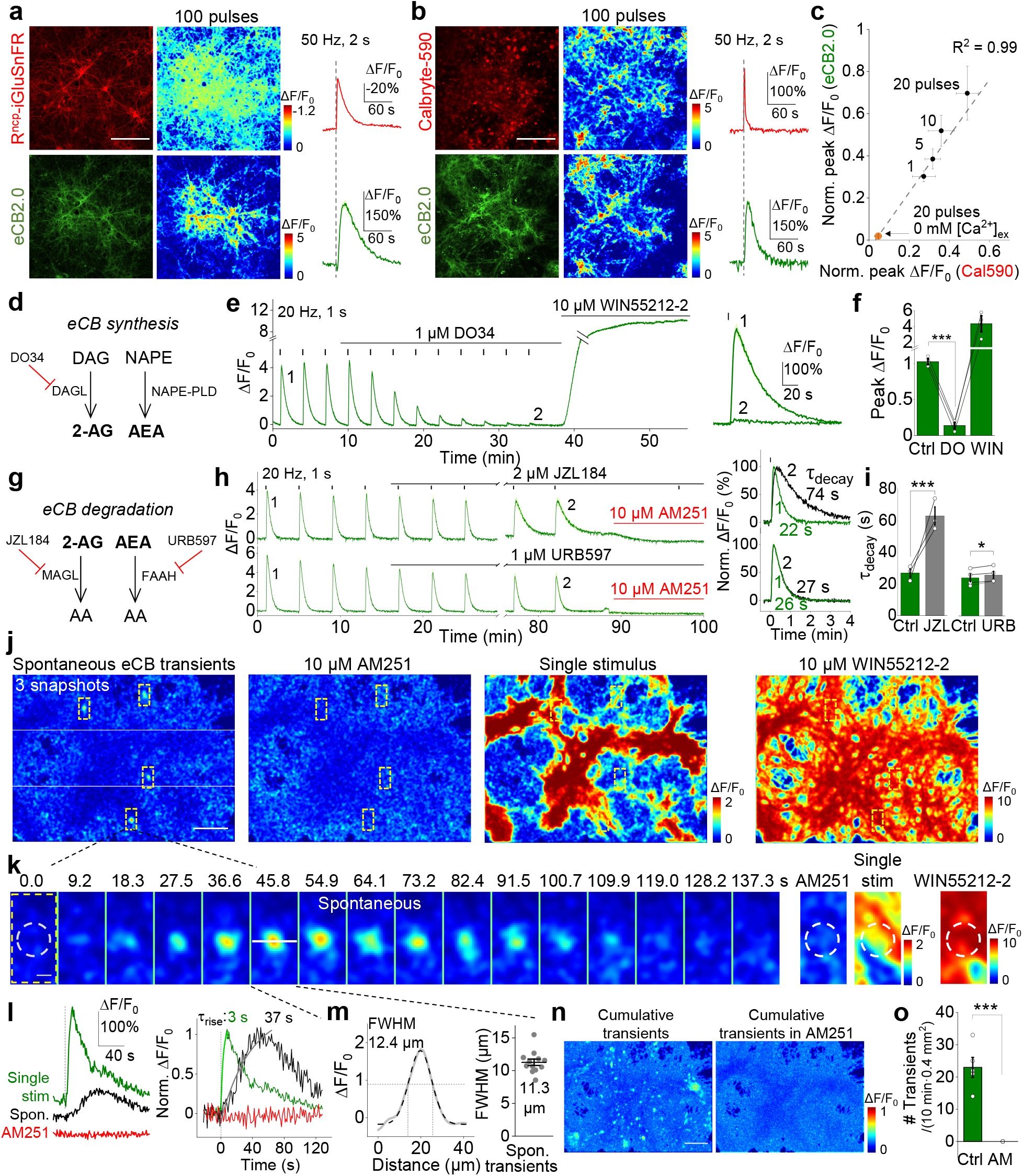
Release of endogenous eCB measured in primary cultured neurons. **a**, Fluorescence microscopy images and fluorescence response measured in neurons co-expressing R^ncp^-iGluSnFR (red) and eCB2.0 (green). Scale bar, 200 μm. **b**, Fluorescence microscopy images and fluorescence response measured in eCB2.0-expressing cells preloaded with Calbryte-590 (red). Scale bar, 200 μm. **c**, Relative peak change in eCB2.0 fluorescence plotted against the relative peak change in Calbryte590 fluorescence measured in response to the indicated number of electrical pulses, normalized to the response evoked by 200 pulses; n = 4 cultures each. Also shown is the response to 20 electrical pulses with no extracellular Ca^2+^. **d**, Diagram depicting the pathway for eCB synthesis. DAG, diacylglycerol; DAGL, diacylglycerol lipase; NAPE, *N*-arachidonoyl phosphatidylethanolamine; NAPE-PLD, NAPE-hydrolyzing phospholipase D. **e**, Representative traces (left) and expanded traces (right) showing the change in eCB2.0 fluorescence in responses to 20 electrical pulses applied before (1) and after (2) DO34 application; WIN55212-2 was applied at the end of the experiment. **f**, Summary of the peak change in eCB2.0 fluorescence in response to 20 pulses applied at baseline (Ctrl), 26 min after DO34 application, and after WIN55212-2 application; n = 3 cultures each. **g**, Diagram depicting the degradation pathways for 2-AG and AEA. AA, arachidonic acid; MAGL, monoacylglycerol lipase; FAAH, fatty acid amide hydrolase. **h**, Representative traces (left) and expanded traces (right) showing the change in eCB2.0 fluorescence in response to 20 electrical pulses applied before (1) and after (2) JZL184 or URB597 application; AM251 was applied at the end of the experiment. **i**, Summary of the decay time constant (Tdecay) measured at baseline (Ctrl) and 68 min after application of either JZL184 or URB597; n = 3 cultures each. **j**, Pseudocolor images showing spontaneous changes in eCB2.0 fluorescence transients, single pulse–evoked fluorescence change, and the change in fluorescence induced by 10 μM WIN55212-2 (note the difference in scale). Scale bar, 100 μm. **k**, Time-lapse pseudocolor images taken from the area shown by the bottom dashed rectangle in panel **j**. Scale bar, 10 μm. **l**, Traces from the experiment shown in panel **k**, showing the change in fluorescence measured spontaneously, induced by a single pulse, or in the presence of AM251. Normalized traces with the corresponding rise time constants are shown at the right. **m**, Spatial profile of the transient change in fluorescence shown in panel **k**. The summary data are shown at the right; n = 12 transients. **n**, Cumulative transient change in eCB2.0 fluorescence measured during 19 mins of recording in the absence (left) or presence (right) of AM251 (right). Pseudocolor images were calculated as the average temporal projection subtracted from the maximum temporal projection. Scale bar, 100 μm. **o**, Summary of the frequency of transient changes in eCB2.0 fluorescence measured before (Ctrl) and after AM251 application; n = 5 & 3 with 10-min recording/session. Student’s *t* tests were performed in **f**, **I** and **o**: *p < 0.05, ***p < 0.001.

Next, we asked which specific eCB—2-AG and/or AEA—is released in cultured rat cortical neurons. 2-AG is mainly produced in neurons from diacylglycerol (DAG) by diacylglycerol lipase (DAGL), while AEA is mainly produced from *N*-arachidonoyl phosphatidylethanolamine (NAPE) via the enzyme NAPE-hydrolyzing phospholipase D (NAPE-PLD) (**Fig. 3d**). We found that the selective DAGL inhibitor DO34^52^ eliminated the stimulus-evoked eCB2.0 signal within 30 min; as a positive control, subsequent application of the CB1R agonist WIN55212-2 restored eCB2.0 fluorescence, indicating that the sensor is still present in the cell membrane (**Fig. 3e,f**). We also examined the effect of blocking the degradation of 2-AG and AEA via the enzymes monoacylglycerol lipase (MAGL) and fatty acid amide hydrolase (FAAH) using the inhibitors JZL184^53^ and URB597^54^, respectively (**Fig. 3g**). We found that blocking MAGL significantly increased the decay time constant, while blocking FAAH had only a slight—albeit significant—effect on the decay time constant. Taken together, these data indicate that 2-AG is the principal eCB released from cultured rat cortical neurons in response to electrical stimuli.

In addition to the stimuli-evoked eCB signals, we also observed local, transient eCB2.0 signals in neurons that occurred spontaneously in the absence of external stimulation (**Fig. 3j**). The peak amplitude and rise kinetics of these transient eCB2.0 signals were smaller and slower compared to the signal measured in response to a single electrical stimulus recording in the same region of interest (ROI) (**Fig. 3k,l**), suggesting that evoked and spontaneous eCB release have distinct patterns. The average diameter of the spontaneous transient signals was 11.3 μm based on our analysis of full width at half maximum (FWHM) (**Fig. 3m**), consistent with previous suggestions that eCB acts locally^55,56^. Finally, the CB1R inverse agonist AM251 eliminated the spontaneous transient eCB2.0 signals (**Fig. 3l,n,o**).

### eCB2.0 can be used to measure eCB release in acute mouse brain slices

Next, we examined whether our eCB sensor can be used to detect endogenous eCB release in a more physiologically relevant system, namely acute mouse brain slices. We first injected AAVs expressing either eCB2.0 or eCBmut into the dorsolateral striatum (DLS) of adult mice (**Fig. 4a**), the region where eCB mediates both short-term and long-term depression and regulates motor behavior^57–59^. Four weeks after AAV injection, acute brain slices were prepared, showing the expression of eCB sensors in DLS (**Fig. 4b**). The fluorescence signals evoked by electrical stimuli in the DLS were recorded by photometry. We found that applying electrical stimuli in eCB2.0-expressing slices evoked clear fluorescence signals, with stronger responses evoked by increasing the number of stimuli and by increasing the stimulation frequency (**Fig. 4c,d**). The half-rise time and decay time constant ranged from 0.8–1.2 s and 5.2–8.5 s, respectively, depending on the number of pulses and the stimulation frequency (**Fig. 4d**). Moreover, the signal was specific to eCB release, as pretreating the slices with 10 μM AM251 blocked the response, and no response was measured in slices expressing the eCBmut mutant sensor (**Fig. 4e**). In a separate experiment, the expression of eCB2.0 in neurites in a striatal slice was detected by 2-photon (2P) fluorescence microscopy; applying AEA induced an increase of eCB2.0 fluorescence that was reversed by AM251 (**Extended Data Fig. 4**).

**Fig. 4.**
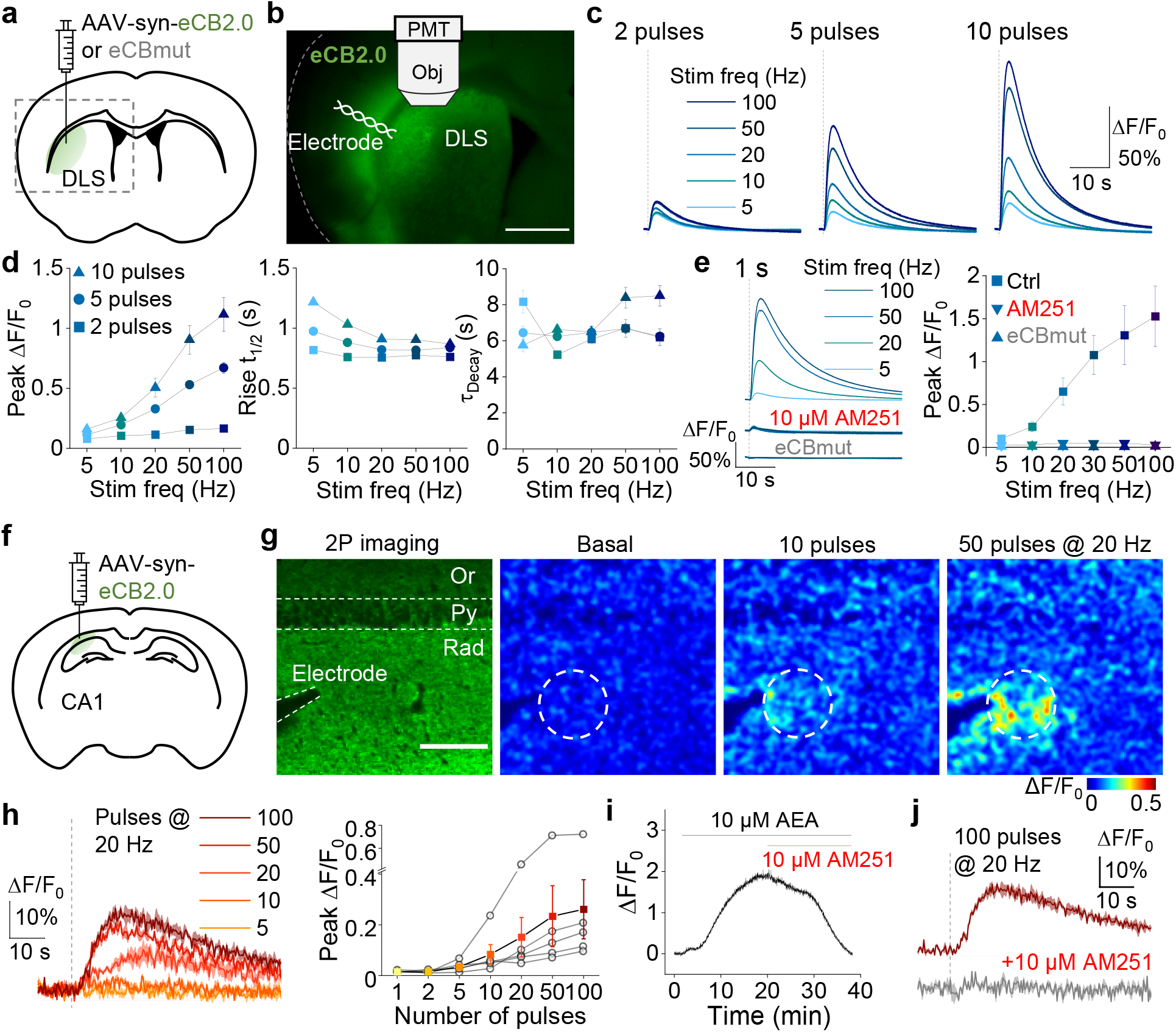
Using the GRAB_eCB_ sensor to detect eCB release in acute brain slices. **a**, Schematic diagram depicting the strategy for virus injection in the dorsolateral striatum (DLS), followed by the preparation of acute brain slices used for electrical stimulation and photometry recording. The dashed box corresponds to the image shown in panel **b**. **b**, Fluorescence image of a coronal slice prepared from a mouse following injection of AAV-syn-eCB2.0 in the DLS, with a diagram showing the electrode position and photometry recording. Scale bar, 1 mm. **c**, Representative traces showing the change in eCB2.0 fluorescence evoked by 2, 5, or 10 electrical pulses applied at the indicated frequencies. **d**, Peak change in eCB2.0 fluorescence (left), rise t_1/2_ (middle), and decay time constant (right) plotted against stimulation frequency for 2, 5, and 10 pulses; n = 6 slices. **e**, Representative traces (left) and summary of the peak change in eCB2.0 fluorescence (right) evoked by electrical pulses at the indicated frequency in slices expressing eCB2.0 in the absence or presence of AM251 and in slices expressing eCBmut; n = 3–4 slices each. **f**, Schematic diagram depicting the strategy for virus injection in the hippocampal CA1 region, followed by the preparation of acute slices for electrical stimulation and 2-photon imaging. **g**, (Left) fluorescence image of eCB2.0 expressed in the hippocampal CA1 region, showing the position of the stimulating electrode. (Right) pseudocolor images showing the change in eCB2.0 fluorescence at baseline and after 10 or 50 pulses applied at 20 Hz. The dashed circle shows the ROI for quantification. Scale bar, 100 μm. **h**, Representative traces and summary of the peak change in eCB2.0 fluorescence evoked by electrical pulses applied at the indicated frequencies; n = 5 slices. **i**, Time course of the change in eCB2.0 fluorescence; where indicated, AEA and AM251 were applied. **j**, Representative traces of the change in eCB2.0 fluorescence evoked by electrical stimulation in the absence and presence of AM251.

We also expressed the eCB2.0 in the hippocampal CA1 region (**Fig. 4f**), in which eCB modulates both excitatory and inhibitory inputs^60,61^, and then recorded eCB2.0 signals in acute slices using 2P microscopy. Consistent with our results measured in in the DLS, we found that applying an increasing number of electrical stimuli at 20 Hz evoked increasingly larger changes in eCB2.0 fluorescence (**Fig. 4g,h**). In addition, applying 10 μM AEA to the slices caused a large increase in eCB2.0 fluorescence that was reversed by 10 μM AM251 (**Fig. 4i**). Finally, AM251 eliminated the signal induced by even 100 field stimuli (**Fig. 4j**). These *in vitro* data confirm that eCB2.0 can be used to reliably detect the endogenous release of eCBs in acute brain slices with high sensitivity, specificity, and spatiotemporal resolution.

### eCB2.0 can be used to measure foot shock-induced eCB release in the basolateral amygdala of freely moving mice

The basolateral amygdala (BLA) is a key brain region mediating fear responses and processing aversive memories^62^. Previous studies found that the CB1R is highly expressed in the BLA, and the eCB system in BLA participates in stress expression^63–65^. We therefore tested whether our eCB2.0 sensor could be used to directly measure eCB dynamics *in vivo* while applying an aversive stimulus (foot shock); for these experiments, we injected AAV vectors expressing either eCB2.0 or eCBmut together with AAVs expressing the mCherry in the mouse BLA and then performed fiber photometry recording (**Fig. 5a,b**). We found that applying a 2-sec foot shock induced a time-locked transient increase in eCB2.0 fluorescence in the BLA (**Fig. 5c**); this response was highly reproducible over 5 consecutive trials (**Fig. 5d**). Importantly, the same foot shock had no effect on either mCherry fluorescence or eCBmut fluorescence (**Fig. 5c,e**). The average time constant for the rise and decay phases of the eCB2.0 signal was 1.0 s and 6.3 s, respectively (**Fig. 5f**). These data indicate that eCB2.0 can be used to measure eCB dynamics *in vivo* in freely moving animals.

**Fig. 5.**
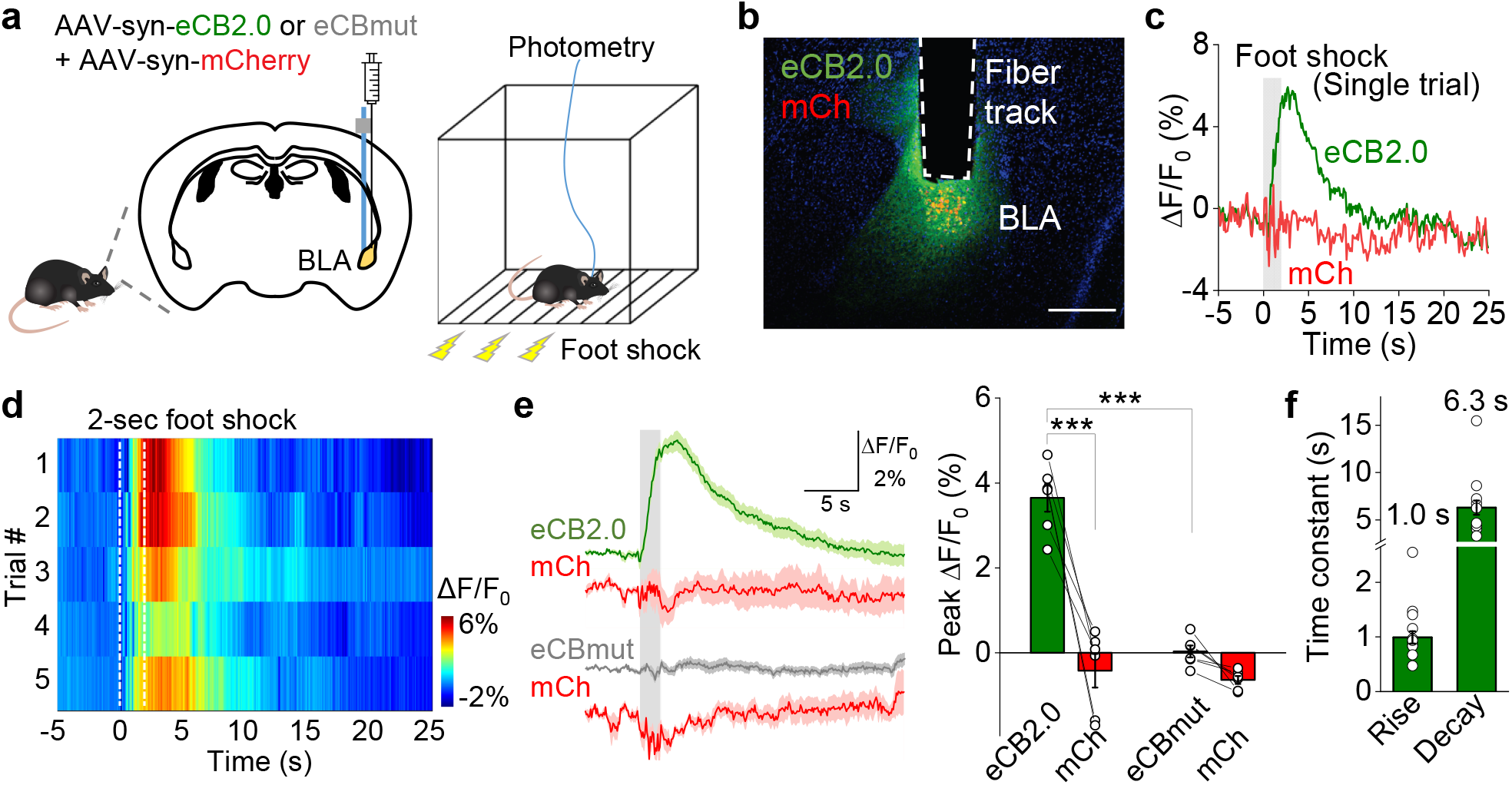
Measuring *in vivo* eCB signals in the mouse basolateral amygdala in response to foot shock. **a**, Schematic diagram depicting the strategy for viral expression in the basolateral amygdala and fiber photometry recording during foot shock. **b**, Fluorescence microscopy image showing eCB2.0 (green) and mCherry (red) expressed in the BLA and the placement of the recording fiber; the nuclei were counterstained with DAPI (blue). Scale bar, 300 μm. **c**, Representative single-trial traces of the change in eCB2.0 and mCherry fluorescence; an electrical foot shock (2-sec duration) was applied at time 0. **d**, Pseudocolor change in eCB2.0 fluorescence before and after a 2-sec foot shock. Shown are five consecutive trials in one mouse, time-aligned to the onset of each foot shock. **e**, (Left) average traces of the change in eCB2.0 and mCherry (top) and eCBmut and mCherry (bottom) fluorescence; the gray shaded area indicates application of an electrical foot shock. (Right) summary of the peak change in fluorescence; n = 6 mice each. **f**, Summary of rise and decay time constants measured for the change in eCB2.0 fluorescence in response to foot shock; n = 18–21 trials in 6 animals. Student’s *t* tests were performed in **e**; ***p < 0.001.

### Dual-color imaging of eCB2.0 and a genetically encoded Ca^2+^ indicator expressed in the mouse hippocampal CA1 region measured during running and seizure activity

Our finding that eCB2.0 can be expressed in the mouse hippocampal CA1 region and then measured in acute slices led us to ask whether we could use this sensor to measure *in vivo* eCB dynamics in the CA1 region during physiologically relevant activity such as running. We therefore injected AAVs expressing eCB2.0 or eCBmut together with a red Ca^2+^ indicator jRGECO1a^66^ into mouse hippocampal CA1 region and then conducted head-fixed 2P dual-color imaging through an implanted cannula above the hippocampus (**Fig. 6a**). Co-expression of eCB2.0 and jRGECO1a was clearly observed in neurons in the CA1 4–6 weeks after virus injection (**Fig. 6b**). We focused on the *stratum pyramidale* layer, which is composed of pyramidal neuron somata and interneuron axons, including a class that densely express CB1R. When mice spontaneously ran on a treadmill (**Fig. 6c**), we found rapid increases of both calcium and eCB signals aligned to the start of running, and decreases of both signals when the running stopped (**Fig. 6d,e**). In the control group, which expressed eCBmut and jRGECO1a, calcium signals were intact while eCBmut showed no fluorescence change (**Fig. 6d,e**). Interestingly, the calcium signal appeared earlier than the eCB signal, although both signals had similar 10%-90% rise time, while the half-time of the decay phase of eCB signal was slower than that of the calcium signal (**Fig. 6f**).

**Fig. 6.**
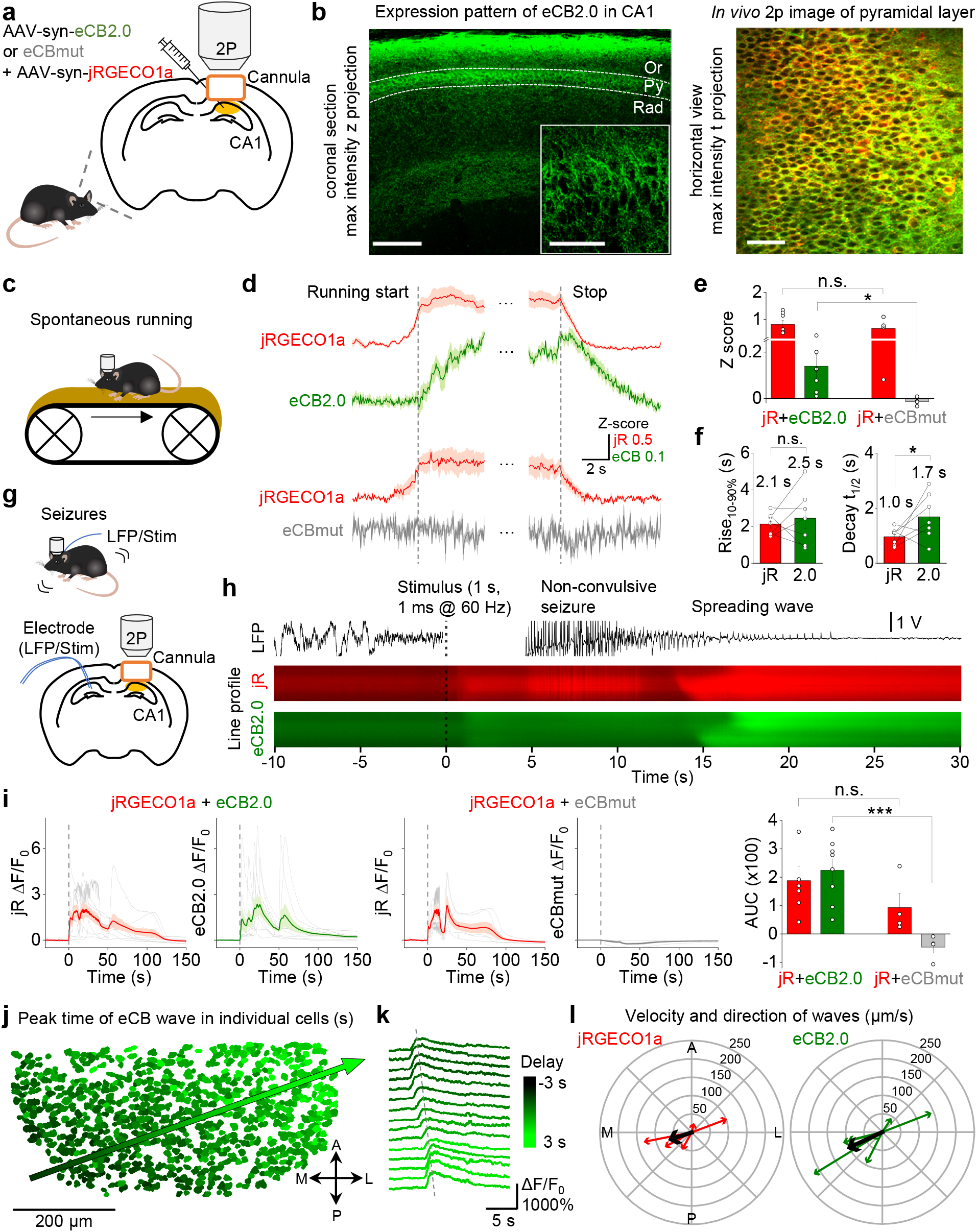
Measuring *in vivo* eCB dynamics in the mouse hippocampus during running and seizure activity. **a**, Schematic diagram depicting the strategy for viral expression and cannula placement in the mouse hippocampus. **b**, (Left) immunofluorescence image showing eCB2.0 expression in the hippocampal CA1 region in a coronal brain slice. Scale bars, 200 μm and 50 μm (inset). (Right) *In vivo* 2-photon image of the pyramidal layer in the hippocampal CA1 region, showing eCB2.0 (green) and jRGECO1a (red) fluorescence. Scale bar, 50 μm. **c**, Schematic cartoon illustrating the experiment in which a mouse expressing eCB2.0 and jRGECO1a in the hippocampal CA1 is placed on a treadmill and allowed to run spontaneously while fluorescence is measured using 2-photon microscopy. **d**, Average traces of eCB2.0/eCBmut and jRGECO1a transients recorded in the soma of individual neurons in the pyramidal layer upon the start and stop of spontaneous running episodes (dashed lines). **e**, Summary of the peak responses in panel **d**; n = 8 and 4 mice each for eCB2.0 and eCBmut, respectively. **f**, Summary of the rise and decay kinetics of the jRGECO1a and eCB2.0 signals measured at the start and end of spontaneous running; n = 7 mice. **g**, Schematic diagram depicting the electrode placement and 2-photon imaging in mice expressing eCB2.0 and jRGECO1a in the hippocampal CA1 region; the electrode is used to induce kindling seizure activity and to measure the local field potential (LFP). **h**, Example LFP trace (top) and medio-lateral projections (line profile) of jRGECO1a (middle) and eCB2.0 (bottom) fluorescence during stimulus-induced non-convulsive seizures and a subsequent spreading wave. The dashed vertical line at time 0 indicates the stimulus onset. **i**, Individual (thin gray lines) and average (thick lines) traces of the change in jRGECO1a and eCB2.0/eCBmut fluorescence measured during seizure activity. The dashed vertical line at time 0 indicates the stimulus onset. The summary of the area under the curve (AUC) is shown at the right; n = 8 and 4 for eCB2.0 and eCBmut, respectively. **j**, Spreading eCB wave measured through the hippocampal CA1 region after seizure activity. ROIs representing individual neurons are pseudocolored based on the peak time of their eCB2.0 signal relative to the peak time of the average signal, and the arrow shows the direction of the wave. a, anterior; l, lateral; m, medial; p, posterior. **k**, Traces of eCB2.0 fluorescence measured in individual cells sampled systematically along a line fitted to the spreading wave. The dashed line shows the spreading of peak signals. **l**, Velocity and direction of the spreading jRGECO1a and eCB2.0 waves. The length of each arrow indicates the velocity in μm/s. In each panel, each colored arrow indicates an individual session, and the thick black line indicates the average. n = 7 sessions in 6 mice. Student’s *t* tests were performed in **e**, **f** and **i**: *p < 0.05, ***p < 0.001, and n.s., not significant.

Epilepsy is a neurological disease characterized by excessive and synchronous neuronal firing. eCBs are proposed to provide negative feedback during epilepsy to attenuate the synaptic activity and protect the nervous system, which is exemplified by the observation that animals with compromised eCB system all exhibit a pro-epileptic phenotype^67^. To explore whether our eCB2.0 sensor could be used to study seizure-related eCB signals *in vivo*, we used electrical kindling stimulation of the hippocampus contralateral to the sensor expressing hemisphere to elicit brief self-terminating seizures (measured using local field potential (LFP) recording) (**Fig. 6g**). We found strong calcium and eCB signal increases during electrical seizure activity (**Fig. 6h**). Recent work has shown that seizures are often followed by a spreading calcium wave that propagates across the cell layer^68^. Interestingly, we also found a propagating eCB wave that closely followed the calcium wave (**Fig. 6h, Extended Data Fig.5 and Supplementary Video 1**). In contrast, eCBmut showed no response during and after seizures (**Fig. 6i**). The velocity and direction of eCB waves were evident when we extracted the eCB2.0 signal from individual neurons in the field of view (**Fig. 6j,k**). Notably, eCB waves and calcium waves varied across experiment sessions and animals (**Fig. 6l**), but for each instance, the calcium and eCB waves were similar, in agreement with the calcium- and activity-dependence of the eCB signal. Taken together, our results confirm that the eCB2.0 sensor can be used to measure eCB dynamics *in vivo* under both physiological and pathological conditions, with high specificity and spatiotemporal resolution.

## DISCUSSION

Here, we report the development and characterization of a genetically-encoded fluorescent sensor for detecting eCBs both *in vitro* and *in vivo*. With high sensitivity, selectivity and kinetics, this novel eCB sensor can be used to detect endogenous eCB release in cultured neurons, acute brain slices and in specific brain structures *in vivo* such as the amygdala and hippocampus during both physiological and pathological activities.

Our estimate of t_on_ and t_off_ kinetics measured for eCB2.0 in cultured neurons at room temperature is likely high, given that a faster time constant was measured in acute slices and in our *in vivo* experiments. Nevertheless, given that the temporal resolution of eCB2.0 is on the order of seconds, this tool is a vast improvement compared to microdialysis (with temporal resolution on the order of minutes), although the sensor’s kinetics could be improved even further in order to capture more rapid signals^69^. In addition, the eCB2.0 sensor can detect both 2-AG and AEA; given that 2-AG and AEA regulate distinct pathways and are involved in different brain regions and cell types^4^, next-generation GRAB_eCB_ sensors should be developed with non-overlapping eCB specificity, as well as non-overlapping color spectra.

The retrograde modulation of synaptic activity by eCBs was previously identified by studying depolarization-induced suppression of inhibition (DSI) and excitation (DSE) in the hippocampus and cerebellum^24,25,27^. However, because these experiments and subsequent studies used electrophysiological recordings of synaptic transmission combined with either pharmacological interventions (e.g., to activate or inhibit eCB receptors or to inhibit enzymes involved in the production or degradation of eCBs) or genetic manipulation (e.g., by knocking out the corresponding receptors and enzymes), they lacked the ability to directly measure eCB release. Moreover, recording at the cell body of a neuron does not provide precise spatial information with respect to eCB release. For example, DSI recorded using paired whole-cell recordings in hippocampal slices indicates that depolarization of one neuron can inhibit GABAergic input to neurons within approximately 20 μm, suggesting the upper limit of diffusion for eCBs from a single neuron^24^; similar results were obtained in cerebellar slices using two separate stimulating electrodes to evoke eCB release from two dendritic regions in a single Purkinje cell^55^. Although these data indicate that eCB signaling is relatively localized and tightly controlled, the detailed spatial profile of eCB signaling is unknown. In addition, although the sampling rate of electrophysiological recordings is generally high (e.g., on the order of several kHz), the eCB signals measured by changes in evoked postsynaptic currents (ePSCs) have a sampling interval of approximately 2 s, creating a temporal bottleneck. In this respect, our eCB2.0 sensor can reveal eCB signals with considerably higher spatial and temporal resolution, similar to recent studies using sensors for detecting other neurotransmitters^70,71^. Using cultured neurons, we found that spontaneous eCB transients are confined to an area with a diameter of approximately 11 μm, smaller than previous estimates of eCB diffusion. In the future, it will be interesting to determine whether these local transient signals originate from single spines.

In summary, we show that our eCB2.0 sensor can be used in a variety of *in vitro* and *in vivo* preparations in order to monitor eCB dynamics in real time. Given the complexity of the nervous system, future directions for research based on the eCB sensor applications may include the identity of cell types that release eCBs, the mechanisms and temporal properties of eCB release, characteristics of eCB diffusion, the duration of eCB signals, the nature of the cell types and subcellular elements targeted by eCBs and the effects on them. Answering these fundamental questions will significantly enrich our understanding of the mechanisms and functions of eCB signaling at the synapse and neural circuit levels. Lastly, altered function of the eCB system has been associated with several neurological disorders, including stress/anxiety, movement disorders, substance use disorders and epilepsy. In this respect, our *in vivo* results show clear examples of how the eCB2.0 sensor could help to elucidate the fast eCB dynamics during both physiological and pathological processes. The eCB2.0 sensor should be able to detect all CB1R agonists (**Extended Data Fig. 3**) including Δ-9-tetrahydrocannabinol (Δ-9-THC) in the brain and periphery following drug administration. This would also allow investigators to track the time course of Δ-9-THC actions and the impact of cannabis drugs on eCB signaling. Thus, eCB sensors open a new era of endocannabinoid research aimed at understanding this system at unprecedented, physiologically-relevant spatial and temporal scales.

## METHODS

### Molecular biology

DNA fragments were amplified by PCR using primers (TSINGKE Biological Technology) with 25–30-bp overlaps. Plasmids were constructed using restriction enzyme cloning or Gibson Assembly, and all plasmid sequences were verified using Sanger sequencing. To characterize eCB2.0 and eCBmut in HEK293T cells, the corresponding DNA constructs were cloned into the pDisplay vector with an upstream IgK leader sequence. An IRES-mCherry-CAAX cassette was inserted downstream of the sensor gene for labeling the cell membrane and calibrating the sensor’s fluorescence. To characterize eCB2.0 in neurons, the eCB2.0 was cloned into a pAAV vector under control of a human synapsin (*SYN1*) promoter (pAAV-hSyn), and PSD95-mScarlet and synaptophysin-mScarlet were cloned into the pDest vector under the control of the *CMV* promoter. For the G_βγ_ sensor assay, the human CB1R was cloned into the pCI vector (Promega), and eCB2.0 and eCBmut were cloned into the peGFP-C1 vector (Takara), replacing the eGFP open reading frame. For the Tango assay, the human CB1R, eCB2.0 and eCBmut were cloned into the pTango vector. In addition, the viral vectors pAAV-hsyn-eCBmut and pAAV-hsyn-R^ncp^-iGluSnFR were generated and used in this study.

### AAV expression

AAV2/9-hSyn-eCB2.0 (9.5×10^13^ viral genomes (vg)/mL), AAV2/9-hSyn-eCBmut (8.0×10^13^ vg/mL), AAV2/9-hSyn-R^ncp^-iGluSniFR (6.2×10^13^ vg/mL, all packaged at Vigene Biosciences, China), AAV8-hSyn-mCherry (#114472, Addgene) and AAV1-Syn-NES-jRGECO1a-WPRE-SV40 (Penn Vector Core) were used to infect cultured neurons or were injected *in vivo* into specific brain regions.

### Cell culture

HEK293T cells were cultured at 37°C in air containing 5% CO_2_ in DMEM (Biological Industries) supplemented with 10% (v/v) fetal bovine serum (Gibco) and penicillin (100 unit/mL)-streptomycin (0.1 mg/mL) (Biological Industries). For experiments, the HEK293T cells were plated on 96-well plates or 12 mm glass coverslips in 24-well plates. At 60–70% confluency, the cells were transfected using polyethylenimine (PEI) with 300 ng DNA/well (for 96-well plates) or 1 μg DNA/well (for 24-well plates) at a DNA:PEI ratio of 1:3; 4–6 h after transfection, the culture medium was replaced with fresh medium. Imaging was performed 24–36 h after transfection. Rat cortical neurons were prepared from postnatal day 0 (P0) Sprague-Dawley rat. In brief, the cerebral cortex was dissected, and cortical neurons were dissociated by digestion in 0.25% Trypsin-EDTA (Biological Industries), and then plated on poly-D-lysine–coated (Sigma-Aldrich) 12-mm glass coverslips in 24-well plates. The neurons were cultured at 37°C, 5% CO_2_ in Neurobasal Medium (Gibco) supplemented with 2% B-27 Supplement (Gibco), 1% GlutaMAX (Gibco), and penicillin (100 unit/mL)-streptomycin (0.1 mg/mL) (Biological Industries). For transfection, cultured neurons were transfected at 7–9 day *in vitro* (DIV7–9) using calcium phosphate transfection method and imaged 48 h after transfection. For viral infection, cultured neurons were infected by AAVs expressing eCB2.0, eCBmut and/or R^ncp^-iGluSnFR at DIV3–5 and imaged at DIV12–20. Where indicated, the neurons were loaded with Calbryte-590 (AAT Bioquest) 1 h before imaging.

### Animals

All experiment protocols were approved by the respective Laboratory Animal Care and Use Committees of Peking University, the National Institute on Alcohol Abuse and Alcoholism, the Cold Spring Harbor Laboratory, and Stanford University, and all studies were performed in accordance with the guidelines established by the US National Institutes of Health. Postnatal day 0 (P0) Sprague-Dawley rats (Beijing Vital River Laboratory) of both sexes and P42–P150 C57BL/6J mice (Beijing Vital River Laboratory and The Jackson Laboratory) of both sexes were used in this study. The mice were housed under a normal 12-h light/dark cycle with food and water available *ad libitum*.

### Confocal imaging of cultured cells

Before imaging, the culture medium was replaced with Tyrode’s solution consisting of (in mM): 150 NaCl, 4 KCl, 2 MgCl_2_, 2 CaCl_2_, 10 HEPES, and 10 glucose (pH 7.4). 0 mM [Ca^2+^]_ex_ solution was modified from Tyrode’s solution with 0 mM CaCl_2_ and additional 2 mM EGTA. HEK293T cells in 96-well plates were imaged using an Opera Phenix high-content screening system (PerkinElmer, USA) equipped with a 20x/0.4 NA objective, a 40x/0.6 NA objective, a 40x/1.15 NA water-immersion objective, a 488 nm laser and a 561 nm laser. Green and red fluorescence were collected using a 525/50 nm emission filter and a 600/30 nm emission filter, respectively. Cells in 12 mm coverslips were imaged using a Ti-E A1 confocal microscopy (Nikon, Japan) equipped with a 10x/0.45 NA objective, a 20x/0.75 NA objective, a 40x/1.35 NA oil-immersion objective, a 488 nm laser and a 561 nm laser. Green and red fluorescence were collected using a 525/50 nm emission filter and a 595/50 nm emission filter, respectively. The following compounds were applied by replacing the Tyrode’s solution (for imaging in 96-well plates) or by either bath application or using a custom-made perfusion system (for imaging cells on 12-mm coverslips): 2-AG (Tocris), AEA (Cayman), AM251 (Tocris), LPA (Tocris), S1P (Tocris), ACh (Solarbio), DA (Sigma-Aldrich), GABA (Tocris), Glu (Sigma-Aldrich), Gly (Sigma-Aldrich), NE (Tocris), 5-HT (Tocris), His (Tocris), Epi (Sigma-Aldrich), Ado (Tocris), Tyr (Sigma-Aldrich), WIN55212-2 (Cayman), DO34 (MedChemExpress), JZL184 (Cayman), and URB597 (Cayman). The micropressure application of drugs was controlled by Pneumatic PicoPump PV800 (World Precision Instruments). Cultured neurons were field stimulated using parallel platinum electrodes positioned 1 cm apart; the electrodes were controlled by a Grass S88 stimulator (Grass Instruments), and 1-ms pulses were applied at 80 V. All imaging experiments were performed at room temperature (22–24°C).

### BRET G_βγ_ sensor assay

Plasmids expressing eCB2.0, eCBmut, or CB1R were co-transfected into HEK293T cells together with a single construct expressing human GNAOa, human GNB1 (fused to amino acids 156–239 of Venus), human GNG2 (fused to amino acids 2–155 of Venus), and NanoLuc fused to the amino terminal 112 amino acids of human Phosducin circularly permutated at amino acids 54/55 (Promega). The NanoLuc/Phosducin fusion portion also contains a kRAS membrane targeting sequence at the carboxy terminal end. Templates for assembly were derived from human whole-brain cDNA (Takara) for all cDNAs, except for the hGNB1 and hGNG2 Venus fusions which were a generous gift from Dr. Nevin Lambert (Augusta University). Approximately 24 hours after transfection, the cells were harvested with 10 mM EDTA in phosphate-buffered saline (PBS, pH 7.2), pelleted, and then resuspended in Dulbecco’s modified PBS (Life Technologies) without Ca^2+^ or Mg^2+^. Furimazine (Promega) was then added at a 1/100 dilution to 100 μl of cell suspension in a black 96-well plate, and BRET was measured using a PHERAstar FS plate reader (Berthold) equipped with a Venus BRET cube. The acceptor (Venus) and donor (NanoLuc) signals were measured at 535 nm and 475 nm, respectively, and net BRET was calculated by subtracting the acceptor/donor ratio of a donor-only sample from the acceptor/donor ratio of each sample. Readings were taken before and 3–4 min after application of 20 μM 2-AG (Tocris) to activate CB1R or the eCB sensor.

### Tango assay

Plasmids expressing eCB2.0, eCBmut, or CB1R were transfected into a reporter cell line expressing a β-arrestin2-TEV fusion gene and a tTA-dependent luciferase reporter gene. 24 h after transfection, cells in 6 well plates were collected after trypsin digestion and plated in 96 well plates. AEA was applied at final concentrations ranging from 0.01 nM to 10 μM. 12 h after luciferase expression, Bright-Glo (Fluc Luciferase Assay System, Promega) was added to a final concentration of 5 μM, and luminescence was measured using the VICTOR X5 multi-label plate reader (PerkinElmer).

### Photometry recording in the dorsolateral striatum in acute mouse brain slices

Adult (>10 weeks of age) male C57BL/6J mice were anesthetized with isoflurane, AAV vectors were injected (300 nl at a rate of 50 nl/min) into the dorsolateral striatum at the following coordinates: A/P: +0.75 mm relative to Bregma; M/L: ±2.5 mm relative to Bregma; and D/V: −3.5 mm). After virus injection, the mice received an injection of ketoprofen (5 mg/kg, s.c.), and postoperative care was provided daily until the mice regained their preoperative weight. After a minimum of 4 weeks following AAV injection, the mice were deeply anesthetized with isoflurane, decapitated, and the brains were removed and placed in ice-cold cutting solution containing (in mM): 194 sucrose, 30 NaCl, 4.5 KCl, 26 NaHCO_3_, 1.2 NaH_2_PO_4_, 10 D-glucose, and 1 MgCl_2_ saturated with 5% CO_2_/95% O_2_. Coronal brain slices (250-μm thickness) were prepared and then incubated at 32°C for 60 min in artificial cerebrospinal fluid (ACSF) containing (in mM): 124 NaCl, 4.5 KCl, 26 NaHCO_3_, 1.2 NaH_2_PO_4_, 10 D-glucose, 1 MgCl_2_, and 2 CaCl_2_. After incubation at 32°C, the slices were kept at room temperature until use. Photometry recordings were acquired using an Olympus BX41 upright epifluorescence microscope equipped with a 40x/0.8 NA water-emersion objective and a FITC filter set. Slices were superfused at 2 ml/min with ACSF (29–31°C). A twisted bipolar polyimide-coated stainless-steel stimulating electrode (~200 μm tip separation) was placed in the DLS just medial to the corpus callosum and slightly below the tissue surface in a region with visible eCB2.0 or eCBmut fluorescence. The sensors were excited using either a 470-nm light-emitting diode (LED) (ThorLabs). Photons passing through a 180-μm^2^ aperture positioned just lateral to the stimulating electrode were directed to a model D-104 photomultiplier tube (PMT) (Photon Technology International). The PMT output was amplified (gain: 0.1 μA/V; time constant: 5 ms), filtered at 50 Hz, and digitized at 250 Hz using a Digidata 1550B and Clampex software (Molecular Devices). For each photometry experiment, GRAB_eCB_ was measured as discrete trials repeated every 3 minutes. For each trial, the light exposure duration was 35–45 seconds in order to minimize GRAB_eCB_ photobleaching while capturing the peak response and the majority of the decay phase. To evoke an eCB transient, a train of 200–500-μs electrical pulses (1.0–1.5 mA) was delivered 5 s after initiating GRAB_eCB_ excitation.

### 2-photon imaging in the hippocampus in acute mouse brain slices

Adult (6–8 weeks of age) C57BL/6J mice of both sexes were anesthetized with an intraperitoneal injection of 2,2,2-tribromoethanol (Avertin, 500 mg/kg body weight, Sigma-Aldrich), and AAV vectors were injected (400 nl at a rate of 46 nl/min) into the hippocampal CA1 region using the following coordinates: A/P: −1.8 mm relative to Bregma; M/L: ±1.0 mm relative to Bregma; and D/V: −1.2 mm. After at least 4 weeks following AAV injection, the mice were deeply anesthetized with an intraperitoneal injection of 2,2,2-tribromoethanol, decapitated, and the brains were removed and placed in ice-cold cutting solution containing (in mM): 110 choline-Cl, 2.5 KCl, 0.5 CaCl_2_, 7 MgCl_2_, 1 NaH_2_PO_4_, 1.3 Na ascorbate, 0.6 Na pyruvate, 25 NaHCO_3_, and 25 glucose saturated with 5% CO_2_/95% O_2_. Coronal brain slices (300-μm thickness) were prepared and incubated at 34°C for approximately 40 min in modified ACSF containing (in mM): 125 NaCl, 2.5 KCl, 2 CaCl_2_, 1.3 MgCl_2_, 1 NaH_2_PO_4_, 1.3 Na ascorbate, 0.6 Na pyruvate, 25 NaHCO_3_, and 25 glucose saturated with 5% CO_2_/95% O_2_. Two-photon imaging were performed using an FV1000MPE 2-photon microscope (Olympus) equipped with a 25x/1.05 NA water-immersion objective and a mode–locked Mai Tai Ti:Sapphire laser (Spectra-Physics). The slices were superfused with modified ACSF (32–34°C) at a rate of 4 ml/min. A 920-nm laser was used to excite the eCB2.0 sensor, and fluorescence was collected using a 495–540-nm filter. For electrical stimulation, a bipolar electrode (cat. number WE30031.0A3, MicroProbes for Life Science) was positioned near the stratum radiatum layer in the CA1 region using fluorescence guidance. Fluorescence imaging and electrical stimulation were synchronized using an Arduino board with custom-written software. All images collected during electrical stimulation were recorded at a frame rate of 2.8 fps with a frame size of 256×192 pixels. The stimulation voltage was 4–6 V, and the pulse duration was 1 ms. Drugs were applied to the imaging chamber by perfusion at a flow rate at 4 ml/min.

### Fiber photometry recording of eCB signals in the basolateral amygdala

Adult (10–12 weeks of age) C57BL/6J mice of both sexes anesthetized, and 300 nl of either a 10:1 mixture of AAV-hSyn-eCB2.0 and AAV-hSyn-mCherry or a 10:1 mixture of AAV-hSyn-eCBmut and AAV-hSyn-mCherry was injected using a glass pipette and a Picospritzer III microinjection system (Parker Hannifin) into the right basolateral amygdala using the following coordinates: A/P: −1.78 mm relative to Bregma; M/L −3.30 mm relative to Bregma; and D/V: −4.53 mm. After injection, a 200-μm diameter, 0.37 NA fiber (Inper) was implanted at the same location and secured using resin cement (3M). A head bar was also mounted to the skull using resin cement. At least 14 days after surgery, photometry recording was performed using a commercial photometry system (Neurophotometrics). A patch cord (0.37 NA, Doric Lenses) was attached to the photometry system and to the fiber secured in the mouse brain. A 470-nm LED was used to excite the GRAB_eCB_ sensors, and a 560-nm LED was used to excite mCherry. The average power level of the LED (measured at the output end of the patch cord) was 160 μW and 25 μW for the GRAB_eCB_ sensors and mCherry, respectively. The recording frequency was 10 Hz, and the photometry data were acquired using Bonsai 2.3.1 software.

For the foot shock experiments, the mice were allowed to move freely in a Habitest shock box (Coulbourn Instruments) inside a lighted soundproof behavior box. The FreezeFrame software program was used to apply triggers to the shock generator (Coulbourn Instruments). Five 2-sec pulses of electricity at an intensity of 0.7 mA were delivered to the shock box, with an interval of 90–120 s between trials. After photometry recording, the animals were deeply anesthetized and perfused with PBS followed by 4% paraformaldehyde (PFA) in PBS. The brains were removed, fixed in 4% PFA overnight, and then dehydrated with 30% sucrose in PBS for 24 h. Brain slices were cut using a Leica SM2010R microtome (Leica Biosystems). Floating brain slices were blocked at room temperature for 2 h with a blocking solution containing 5% (w/v) BSA and 0.1% Triton X-100 in PBS, and then incubated at 4°C for 24 h in PBS containing 3% BSA, 0.1% Triton X-100, and the following primary antibodies: chicken anti-GFP (1:1000, Aves, #GFP-1020) and rabbit anti-RFP (1:500, Rockland, #600-401-379). The next day, the slices were rinsed 3 times in PBS and incubated in PBS with DAPI (5 μg/ml, Invitrogen, #D1306) and the following secondary antibodies at 4°C for 24 h: Alexa Fluor 488 donkey anti-chicken (1:250, Jackson ImmunoResearch, #703-545-155) and Alexa Fluor 568 donkey anti-rabbit (1:250, Invitrogen, #A10042). Confocal images were captured using an LSM780 confocal microscope (Zeiss).

### 2-photon *in vivo* imaging

Adult (100–150 days of age) C57BL/6J mice of both sexes were used for these experiments. The mice were anesthetized, and a mixture of AAV1-Syn-NES-jRGECO1a-WPRE-SV40 and either AAV9-hSyn-eCB2.0 or AAV9-hSyn-eCBmut (300–400 nl each, full titer) was injected into the right hippocampal CA1 region at the following coordinates using a Hamilton syringe: A/P: 2.3 mm relative to Bregma; M/L: 1.5 mm relative to Bregma; and D/V: −1.35 mm. After virus injection, a stainless-steel cannula with an attached coverglass was implanted over the hippocampus as described previously^72,73^, and a stainless-steel head bar was attached. A chronic bipolar wire electrode (tungsten, 0.002”, 0.5-mm tip separation, A-M Systems) was implanted into the left ventral hippocampus at the following coordinates as previously described^74^: A/P: 3.2 mm relative to Bregma; M/L: 2.7 mm relative to Bregma; and D/V: −4.0 mm. Head-fixed mice running on a linear treadmill with a 2-m-long cue-less belt were imaged using a resonant scanning 2-photon microscope (Neurolabware) equipped with a pulsed IR laser tunned to 1000 nm (Mai Tai, Spectra-Physics), GaAsP PMT detectors (H11706P-40, Hamamatsu), and a 16x/0.8 NA water-immersion objective (Nikon). The 2-photon image acquisition and treadmill speed were controlled and monitored using a Scanbox (Neurolabware). Bipolar electrodes were recorded using a model 1700 differential amplifier (A-M Systems). Seizures were elicited by applying an electric stimulation above the seizure threshold by 150 μA of current delivered in 1-ms biphasic pulses at 60 Hz for 1 s, using a model 2100 constant-current stimulator (A-M Systems). Following the *in vivo* recordings, the mice were anesthetized with isoflurane followed by an intraperitoneal injection of a mixture of ketamine (100 mg/kg body weight) and xylazine (10 mg/kg body weight) in saline. The mice were transcardially perfused with 0.9% NaCl for 1 min followed by 4% PFA and 0.2% picric acid in 0.1 M phosphate buffer. The brains were removed, post-fixed in the same fixative solution for 24 h at 4°C, then sliced on a VTS1200 vibratome (Leica Biosystems). The sections were then washed and mounted using VECTASHIELD (Vector Laboratories). Confocal images were acquired using an LSM710 imaging system equipped with a 20x/0.8 NA objective (Zeiss).

### Data processing

#### Confocal imaging

Data for 96-well plate imaging were collected and analyzed using Harmony high-content imaging and analysis software (PerkinElmer). In brief, membrane regions were selected as regions of interest (ROIs) and the green fluorescence channel (i.e., the sensor) was normalized to the red fluorescence channel corresponding to mCherry-CAAX (G/R). ΔF/F_0_ was then calculated using the formula [(G/R_drug_ – G/R_baseline_)/(G/R_baseline_)]. For 12-mm coverslip imaging, data were collected using the NIS-Element software (Nikon) and analyzed using ImageJ software (National Institutes of Health). ΔF/F_0_ was calculated as using the formula [(F_t_ – F_0_)/F_0_], with F_0_ representing baseline fluorescence. Data were plotted using OriginPro 2020 (OriginLab).

#### Slice photometry and 2-photon imaging

For slice photometry, GRAB_eCB_ signals were calculated as ΔF/F_0_ by averaging the PMT voltage (V) for a period of 1 s just prior to electrical stimulation (F_0_) and then calculating [V/(F_0_-1)] for each digitized data sample. The decay phase was fitted with a single exponential, accounting for a sloping baseline. Rise t_1/2_ was calculated in Prism v. 8.3(GraphPad) by fitting the rising phase of the signal with an asymmetrical logistics curve. Photometry sweeps were exported to Microsoft Excel 2016 to calculate normalized ΔF/F_0_ traces and peak ΔF/F_0_ values. For 2-photon imaging of slices, data were collected using FV10-ASW software (Olympus) and analyzed using ImageJ. ΔF/F_0_ was calculated using the formula [(F_t_ – F_0_)/F_0_], with F_0_ representing baseline fluorescence. Data were plotted using OriginPro 2020.

#### Fiber photometry in mice during foot shock

The fiber photometry data were analyzed off-line using MatLab software (MathWorks) and plotted using OriginPro 2020.

#### 2-photon imaging in mice during locomotion and seizure

Imaging data were processed and analyzed using Python scripts. To analyze single-cell responses, movies were initially motion-corrected using rigid translation, followed by non-rigid correction (*HiddenMarkov2D*) using the sima package^75^. Binary ROIs were selected using a semi-automated approach. For the initial automated detection, movies were divided into segments consisting of 100 frames each; the average intensity projection of each segment was then computed, and the resulting resampled movie was used for detection. In sessions with electric stimulation, only the baseline period (i.e., before stimulation) was used for segmentation. The *PlaneCA1PC* method of sima was run on the inverted resampled movie, which resulted in detection of the hollow cell nuclei. These ROIs were then filtered based on size, and binary dilation was performed to include the cytoplasm around the nuclei. Next, the ROIs were detected in the non-inverted resampled movie and filtered based on size; those samples that did not overlap with existing ROIs were added to the set. ROIs outside the stratum pyramidale layer were excluded. The fluorescence intensity traces were then extracted for each ROI by averaging the included pixel intensities within each frame. For analyzing the run responses, only sessions with no electric stimuli were included, and signals were pulled from the motion-corrected movies. These raw traces were then processed following standard steps for obtaining ΔF/F_0_ traces, with a modified approach for determining the time-dependent baseline. A 3rd-degree polynomial was fit to the trace after applying temporal smoothing, removing peaks (detected using continuous wavelet transform with scipy.signal), eliminating periods of running, and ignoring the beginning and end of the recording. The calculated polynomial was then used as a baseline. *Z*-scored traces were obtained after determining the standard deviation (SD) of each cell’s baseline and excluding events exceeding 2 SDs in two iterations.

To analyze spreading activity, only sessions with an electric stimulus that triggered an electrographic seizure and a spreading wave were included. The segmentation was performed based on the motion-corrected baseline segments of the recordings, and the signals were pulled from non-motion-corrected movies, as image-based motion correction was not feasible during seizures. ΔF/F_0_ traces were obtained using a constant baseline determined by averaging the pre-stimulus segments of the traces. To analyze changes in average fluorescence intensity, a single large ROI was manually drawn to include the cell bodies within the pyramidal layer, and ΔF/F_0_ traces were obtained and processed as described above. Event-triggered averages were calculated after automatically detecting the frames with running onsets and stops using criteria that were fixed across all sessions. The average was computed in two steps; first, the events were averaged by cell, and then the cells were averaged by sensor (e.g., eCB2.0 or eCBmut). Decay time constants were computed as the parameter of a 2nd-degree polynomial fit after a log transform on the trace following the peak of the stop-triggered average trace. Rise times were determined between the frame in which the start-triggered average signal first reached 90% of the range between baseline and peak and the last frame before the signal dropped below 10% of the range. To determine the speed and direction of the spreading waves, the peak time of the wave was determined in each session by inspecting the average ΔF/F_0_ trace (including all cells). Next, the relative peak location (Δt) of the ΔF/F_0_ trace of each cell in the trace including 200 frames (12.8 s) before and after the wave peak was determined. Finally, two linear (i.e., 1D) fits were determined using the *x* and *y* centroid coordinates of each ROI (Δt ~ x, Δt ~ y). The 2D speed was then computed from the slopes of the two 1D fits. The direction was determined by computing the unity vector from the starting point to the end point of the fits between 3 s before and after the wave peak. The average speed was obtained by averaging the speed of individual sessions, and the average direction was obtained from the sum of the unity vectors of individual sessions. Data were plotted using Python and OriginPro 2020.

### Statistical analysis

All summary data are presented as the mean ± s.e.m. Group data were analyzed using the Student’s *t* test or one-way ANOVA. \**p* < 0.05, \*\**p* < 0.01, \*\*\**p* < 0.001, and n.s., not significant (*p* > 0.05).

### Data and software availability

Plasmids for expressing eCB2.0 and eCBmut used in this study were deposited at Addgene (https://www.addgene.org/Yulong_Li/).

## ACKNOWLEDGMENTS

This work was supported by the Beijing Municipal Science & Technology Commission (Z181100001318002, Z181100001518004), the National Natural Science Foundation of China (31925017), the NIH BRAIN Initiative (NS103558), the Shenzhen-Hong Kong Institute of Brain Science (NYKFKT2019013), and the Peking-Tsinghua Center for Life Sciences and the State Key Laboratory of Membrane Biology at Peking University School of Life Sciences to Y.L.; the NIAAA (ZIA AA000416) to D.M.L; the NIH BRAIN Initiative (NS103558) to J.D.; the NIH (R01MH101214 and R01NS104944) to B.L.; the American Epilepsy Society (postdoctoral fellowship) and the NIH (K99NS117795) to B.D.; the Canadian Institutes for Health Research (postdoctoral fellowship) to J.S.F.; and the NIH to I.S. (NS99457). We thank Li lab members and alumni for helpful discussions. We thank Yi Rao for use of the 2-photon microscope, Xiaoguang Lei at PKU-CLS and the National Center for Protein Sciences at Peking University for support and assistance with the Opera Phenix high-content screening system.

## AUTHOR CONTRIBUTIONS

Y.L. conceived the project. A.D., K.H., H.L.P., R.C., and J.D. performed the experiments related to developing, optimizing, and characterizing the sensors in cultured HEK293T cells and neurons. L.J.D. performed the surgery and photometry recording experiments related to the validation of the sensor in DLS brain slices under the supervision of D.M.L. A.D. performed the surgery and 2-photon imaging in the hippocampal brain slices. E.A. performed the surgery and 2-photon imaging in the striatal brain slices under the supervision of J.D. W. G. performed fiber photometry recordings in freely moving mice during foot shock under the supervision of B.L. B.D. and J.S.F. performed the *in vivo* 2-photon imaging in the hippocampus in mice during running and seizure under the supervision of I.S. All authors contributed to the data interpretation and analysis. A.D. and Y.L. wrote the manuscript with input from other authors.

---

**Extended Data Fig. 1.**
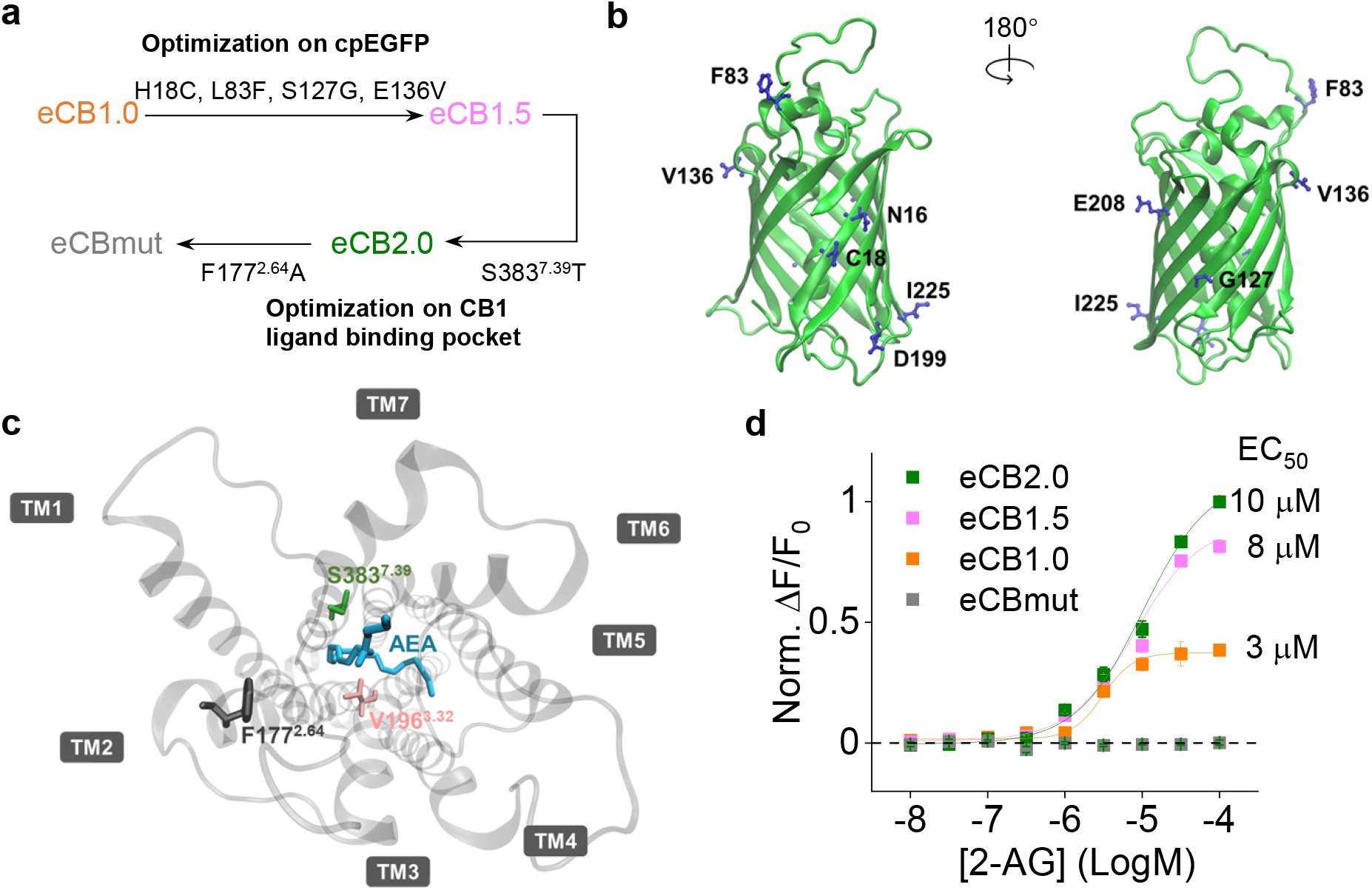
Strategy for optimizing and screening the GRABeCB sensor prototypes. **a**, Schematic diagram depicting the strategy used to generate the various GRAB_eCB_ sensors for this study, including intermediate steps. **b**, Location of the 8 residues in the cpEGFP moiety used to optimize the GRAB_eCB_ sensor. **c**, Location of the 3 residues in the GPCR ligand-binding pocket. The receptor’s seven transmembrane domains (TM1 through TM7) and the ligand molecule (AEA) are shown. **d**, Normalized dose-response curves for the change in eCB1.0, eCB1.5, eCB2.0, and eCBmut fluorescence in response to 2-AG measured in HEK293T cells; n = 3 wells each.

**Extended Data Fig. 2.**
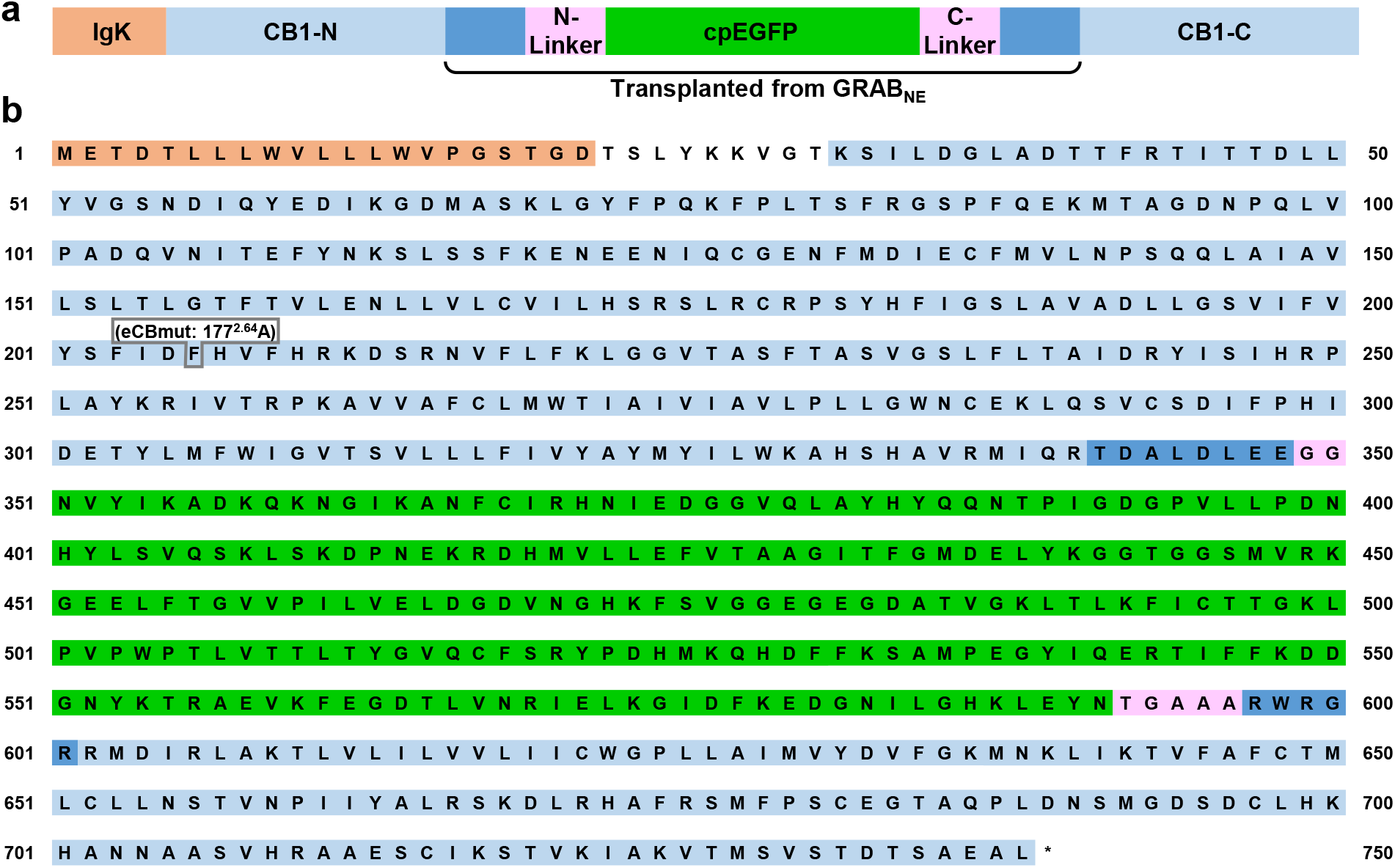
Full amino acid sequences of the eCB2.0 and eCBmut sensors. **a**, Schematic diagram depicting the structure of the GRAB_eCB2.0_ sensor. The IgK leader sequence and the sequence derived from GRAB_NE_ are shown. **b**, Amino acids sequence of the eCB2.0 sensor. The phenylalanine residue at position 177^2.64^ in the CB1 receptor was mutated to an alanine to generate the eCB mutant sensor (indicated by the gray box). Note that the numbering used in the figure corresponds to the start of the IgK leader sequence.

**Extended Data Fig. 3.**
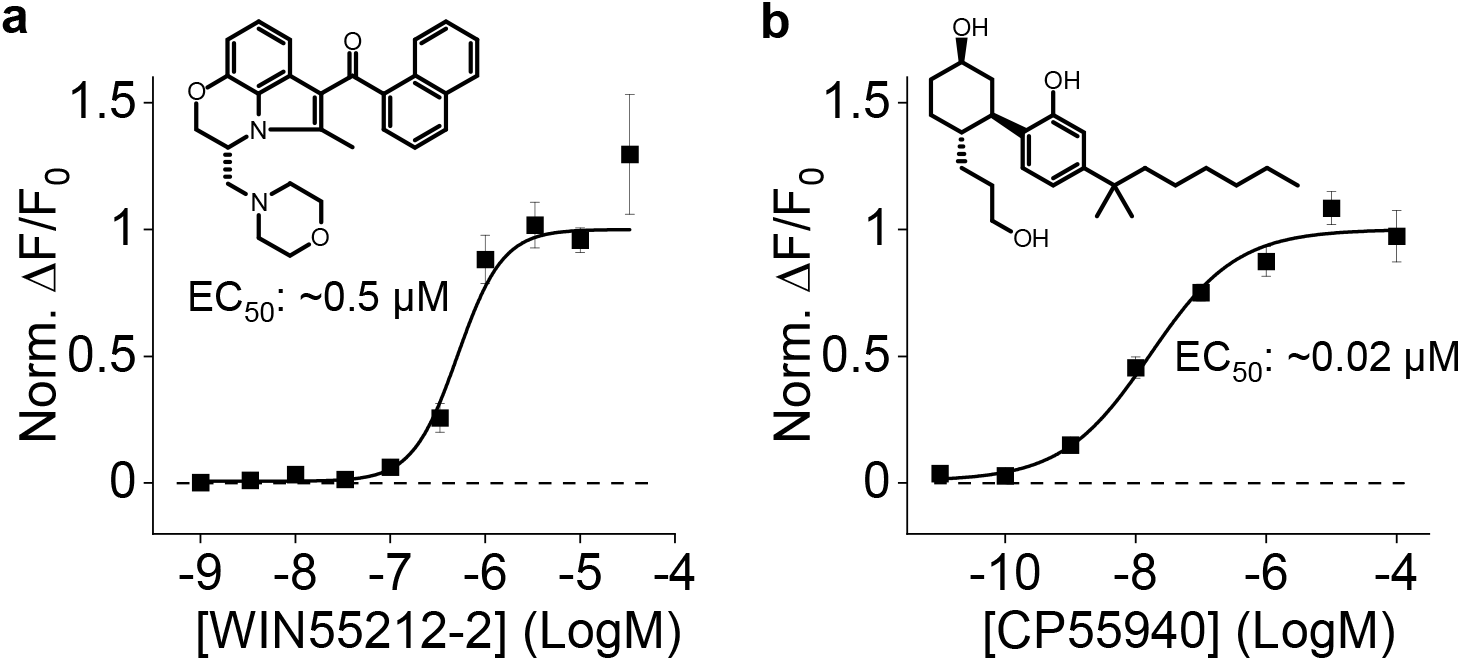
The eCB2.0 responses to synthetic CB1R agonists. **a**, Dose-response curves for WIN55212-2 measured in HEK293T cells expressing eCB2.0, with the corresponding structure and EC_50_ value shown; n = 3 wells each. **b**, Dose-response curves for CP55940 measured in HEK293T cells expressing eCB2.0, with the corresponding structure and EC_50_ value shown; n = 3 wells each.

**Extended Data Fig. 4.**
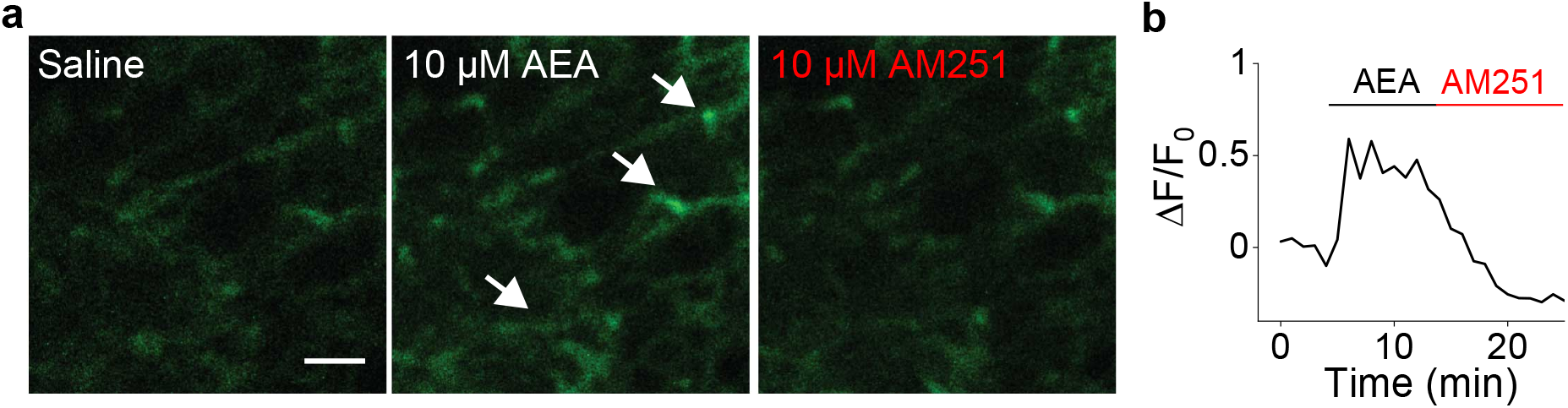
Expression and response of eCB2.0 in acute mouse striatal slices. **a**, Two-photon fluorescence images of eCB2.0 expressed in the striatum before (saline) and after AEA and AM251 application. Arrows indicate eCB2.0 expressing neurites. Scale bar, 10 μm. **b**, Time course of the change in eCB2.0 fluorescence; where indicated, AEA and AM251 were applied.

**Extended Data Fig. 5.**
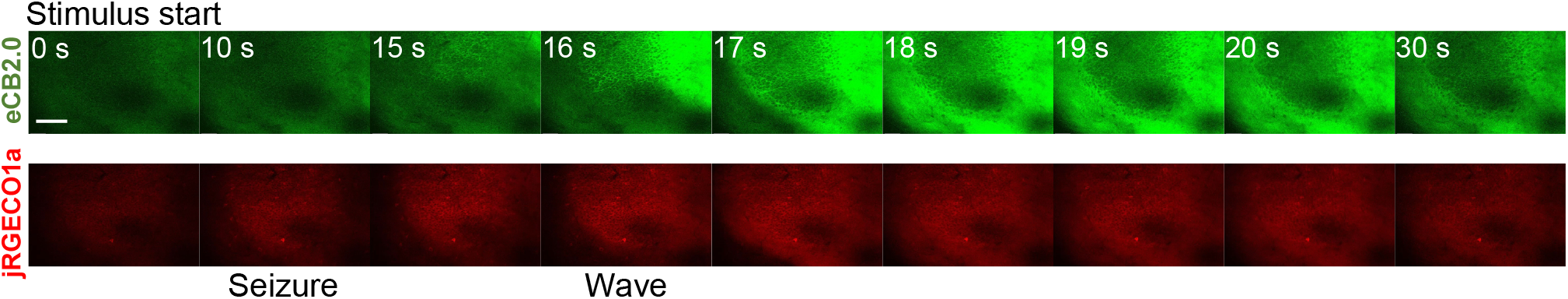
eCB and Ca^2+^ waves in mouse hippocampal CA1 region during seizure activity. *In vivo* two-photon fluorescence images of eCB2.0 and jRGECO1a expressed in the mouse hippocampal CA1 region before and after stimulus evoked seizure activity. Frames were extracted from those shown in Supplementary Video 1. Seconds (s) after the stimulus are indicated. Scale bar, 100 μm.

**Supplementary video 1 | eCB and calcium signals in mouse hippocampal CA1 during seizures**

Fluorescence movies of eCB2.0 and jRGECO1a in the mouse hippocampal CA1 region during seizure activity, which is indicated by the LFP recording. The video is played at 3 times the speed.

## REFERENCES

1 Zuardi, A. W. History of cannabis as a medicine: a review. Braz J Psychiatry 28, 153–157, doi:10.1590/s1516-44462006000200015 (2006).

2 Piomelli, D. The molecular logic of endocannabinoid signalling. Nat Rev Neurosci 4, 873–884, doi:10.1038/nrn1247 (2003).

3 Wilson, R. I. & Nicoll, R. A. Endocannabinoid signaling in the brain. Science 296, 678–682, doi:10.1126/science.1063545 (2002).

4 Kano, M., Ohno-Shosaku, T., Hashimotodani, Y., Uchigashima, M. & Watanabe, M. Endocannabinoid-mediated control of synaptic transmission. Physiological reviews 89, 309–380, doi:10.1152/physrev.00019.2008 (2009).

5 Hebert-Chatelain, E. et al. A cannabinoid link between mitochondria and memory. Nature 539, 555–559, doi:10.1038/nature20127 (2016).

6 Benard, G. et al. Mitochondrial CB(1) receptors regulate neuronal energy metabolism. Nat Neurosci 15, 558–564, doi:10.1038/nn.3053 (2012).

7 Jimenez-Blasco, D. et al. Glucose metabolism links astroglial mitochondria to cannabinoid effects. Nature 583, 603–608, doi:10.1038/s41586-020-2470-y (2020).

8 Stella, N. Cannabinoid signaling in glial cells. Glia 48, 267–277, doi:10.1002/glia.20084 (2004).

9 Navarrete, M., Diez, A. & Araque, A. Astrocytes in endocannabinoid signalling. Philos Trans R Soc Lond B Biol Sci 369, 20130599, doi:10.1098/rstb.2013.0599 (2014).

10 Chevaleyre, V., Takahashi, K. A. & Castillo, P. E. Endocannabinoid-mediated synaptic plasticity in the CNS. Annu Rev Neurosci 29, 37–76, doi:10.1146/annurev.neuro.29.051605.112834 (2006).

11 Oddi, S., Scipioni, L. & Maccarrone, M. Endocannabinoid system and adult neurogenesis: a focused review. Curr Opin Pharmacol 50, 25–32, doi:10.1016/j.coph.2019.11.002 (2020).

12 Moreira, F. A. & Lutz, B. The endocannabinoid system: emotion, learning and addiction. Addict Biol 13, 196–212, doi:10.1111/j.1369-1600.2008.00104.x (2008).

13 Guindon, J. & Hohmann, A. G. The endocannabinoid system and pain. CNS Neurol Disord Drug Targets 8, 403–421 (2009).

14 Kesner, A. J. & Lovinger, D. M. Cannabinoids, Endocannabinoids and Sleep. Frontiers in molecular neuroscience 13, 125, doi:10.3389/fnmol.2020.00125 (2020).

15 Silvestri, C. & Di Marzo, V. The endocannabinoid system in energy homeostasis and the etiopathology of metabolic disorders. Cell Metab 17, 475–490, doi:10.1016/j.cmet.2013.03.001 (2013).

16 Katona, I. & Freund, T. F. Endocannabinoid signaling as a synaptic circuit breaker in neurological disease. Nat Med 14, 923–930, doi:10.1038/nm.f.1869 (2008).

17 Fernandez-Espejo, E., Viveros, M. P., Nunez, L., Ellenbroek, B. A. & Rodriguez de Fonseca, F. Role of cannabis and endocannabinoids in the genesis of schizophrenia. Psychopharmacology (Berl) 206, 531–549, doi:10.1007/s00213-009-1612-6 (2009).

18 Fraguas-Sanchez, A. I., Martin-Sabroso, C. & Torres-Suarez, A. I. Insights into the effects of the endocannabinoid system in cancer: a review. Br J Pharmacol 175, 2566–2580, doi:10.1111/bph.14331 (2018).

19 Patel, S., Hill, M. N., Cheer, J. F., Wotjak, C. T. & Holmes, A. The endocannabinoid system as a target for novel anxiolytic drugs. Neurosci Biobehav Rev 76, 56–66, doi:10.1016/j.neubiorev.2016.12.033 (2017).

20 Ligresti, A., Petrosino, S. & Di Marzo, V. From endocannabinoid profiling to ‘endocannabinoid therapeutics’. Curr Opin Chem Biol 13, 321–331, doi:10.1016/j.cbpa.2009.04.615 (2009).

21 Sabatini, B. L. & Regehr, W. G. Timing of synaptic transmission. Annu Rev Physiol 61, 521–542, doi:10.1146/annurev.physiol.61.1.521 (1999).

22 Zoerner, A. A. et al. Quantification of endocannabinoids in biological systems by chromatography and mass spectrometry: a comprehensive review from an analytical and biological perspective. Biochim Biophys Acta 1811, 706–723, doi:10.1016/j.bbalip.2011.08.004 (2011).

23 Marchioni, C. et al. Recent advances in LC-MS/MS methods to determine endocannabinoids in biological samples: Application in neurodegenerative diseases. Anal Chim Acta 1044, 12–28, doi:10.1016/j.aca.2018.06.016 (2018).

24 Wilson, R. I. & Nicoll, R. A. Endogenous cannabinoids mediate retrograde signalling at hippocampal synapses. Nature 410, 588–592, doi:10.1038/35069076 (2001).

25 Kreitzer, A. C. & Regehr, W. G. Retrograde inhibition of presynaptic calcium influx by endogenous cannabinoids at excitatory synapses onto Purkinje cells. Neuron 29, 717–727 (2001).

26 Maejima, T., Hashimoto, K., Yoshida, T., Aiba, A. & Kano, M. Presynaptic inhibition caused by retrograde signal from metabotropic glutamate to cannabinoid receptors. Neuron 31, 463–475 (2001).

27 Ohno-Shosaku, T., Maejima, T. & Kano, M. Endogenous cannabinoids mediate retrograde signals from depolarized postsynaptic neurons to presynaptic terminals. Neuron 29, 729–738 (2001).

28 Wiskerke, J. et al. Characterization of the effects of reuptake and hydrolysis inhibition on interstitial endocannabinoid levels in the brain: an in vivo microdialysis study. ACS Chem Neurosci 3, 407–417, doi:10.1021/cn300036b (2012).

29 Walker, J. M., Huang, S. M., Strangman, N. M., Tsou, K. & Sanudo-Pena, M. C. Pain modulation by release of the endogenous cannabinoid anandamide. Proc Natl Acad Sci U S A 96, 12198–12203, doi:10.1073/pnas.96.21.12198 (1999).

30 Sun, F. et al. A Genetically Encoded Fluorescent Sensor Enables Rapid and Specific Detection of Dopamine in Flies, Fish, and Mice. Cell 174, 481–496 e419, doi:10.1016/j.cell.2018.06.042 (2018).

31 Patriarchi, T. et al. Ultrafast neuronal imaging of dopamine dynamics with designed genetically encoded sensors. Science 360, doi:10.1126/science.aat4422 (2018).

32 Jing, M. et al. A genetically encoded fluorescent acetylcholine indicator for in vitro and in vivo studies. Nat Biotechnol 36, 726–737, doi:10.1038/nbt.4184 (2018).

33 Feng, J. et al. A Genetically Encoded Fluorescent Sensor for Rapid and Specific In Vivo Detection of Norepinephrine. Neuron 102, 745–761 e748, doi:10.1016/j.neuron.2019.02.037 (2019).

34 Peng, W. et al. Regulation of sleep homeostasis mediator adenosine by basal forebrain glutamatergic neurons. Science 369, doi:10.1126/science.abb0556 (2020).

35 Patriarchi, T. et al. An expanded palette of dopamine sensors for multiplex imaging in vivo. Nat Methods, doi:10.1038/s41592-020-0936-3 (2020).

36 Jing, M. et al. An optimized acetylcholine sensor for monitoring in vivo cholinergic activity. Nature Methods, doi:10.1038/s41592-020-0953-2 (2020).

37 Sun, F. et al. New and improved GRAB fluorescent sensors for monitoring dopaminergic activity <em>in vivo</em>. 2020.2003.2028.013722, doi:10.1101/2020.03.28.013722 %J bioRxiv (2020).

38 Wan, J. et al. A genetically encoded GRAB sensor for measuring serotonin dynamics <em>in vivo</em>. 2020.2002.2024.962282, doi:10.1101/2020.02.24.962282 %J bioRxiv (2020).

39 Howlett, A. C. et al. International Union of Pharmacology. XXVII. Classification of cannabinoid receptors. Pharmacol Rev 54, 161–202, doi:10.1124/pr.54.2.161 (2002).

40 Hua, T. et al. Crystal structures of agonist-bound human cannabinoid receptor CB1. Nature 547, 468–471, doi:10.1038/nature23272 (2017).

41 Hua, T. et al. Crystal Structure of the Human Cannabinoid Receptor CB1. Cell 167, 750–762 e714, doi:10.1016/j.cell.2016.10.004 (2016).

42 Krishna Kumar, K. et al. Structure of a Signaling Cannabinoid Receptor 1-G Protein Complex. Cell 176, 448–458 e412, doi:10.1016/j.cell.2018.11.040 (2019).

43 Li, X. et al. Crystal Structure of the Human Cannabinoid Receptor CB2. Cell 176, 459–467 e413, doi:10.1016/j.cell.2018.12.011 (2019).

44 Shao, Z. et al. Structure of an allosteric modulator bound to the CB1 cannabinoid receptor. Nat Chem Biol 15, 1199–1205, doi:10.1038/s41589-019-0387-2 (2019).

45 Shao, Z. et al. High-resolution crystal structure of the human CB1 cannabinoid receptor. Nature, doi:10.1038/nature20613 (2016).

46 Masuho, I. et al. Distinct profiles of functional discrimination among G proteins determine the actions of G protein-coupled receptors. Sci Signal 8, ra123, doi:10.1126/scisignal.aab4068 (2015).

47 Hollins, B., Kuravi, S., Digby, G. J. & Lambert, N. A. The c-terminus of GRK3 indicates rapid dissociation of G protein heterotrimers. Cell Signal 21, 1015–1021, doi:10.1016/j.cellsig.2009.02.017 (2009).

48 Kroeze, W. K. et al. PRESTO-Tango as an open-source resource for interrogation of the druggable human GPCRome. Nat Struct Mol Biol 22, 362–369, doi:10.1038/nsmb.3014 (2015).

49 Kim, S. H., Won, S. J., Mao, X. O., Jin, K. & Greenberg, D. A. Molecular mechanisms of cannabinoid protection from neuronal excitotoxicity. Mol Pharmacol 69, 691–696, doi:10.1124/mol.105.016428 (2006).

50 Wu, J. et al. Genetically Encoded Glutamate Indicators with Altered Color and Topology. ACS Chem Biol 13, 1832–1837, doi:10.1021/acschembio.7b01085 (2018).

51 Alger, B. E. Retrograde signaling in the regulation of synaptic transmission: focus on endocannabinoids. Prog Neurobiol 68, 247–286, doi:10.1016/s0301-0082(02)00080-1 (2002).

52 Ogasawara, D. et al. Rapid and profound rewiring of brain lipid signaling networks by acute diacylglycerol lipase inhibition. Proc Natl Acad Sci U S A 113, 26–33, doi:10.1073/pnas.1522364112 (2016).

53 Long, J. Z. et al. Selective blockade of 2-arachidonoylglycerol hydrolysis produces cannabinoid behavioral effects. Nat Chem Biol 5, 37–44, doi:10.1038/nchembio.129 (2009).

54 Mor, M. et al. Cyclohexylcarbamic acid 3’-or 4’-substituted biphenyl-3-yl esters as fatty acid amide hydrolase inhibitors: synthesis, quantitative structure-activity relationships, and molecular modeling studies. J Med Chem 47, 4998–5008, doi:10.1021/jm031140x (2004).

55 Brenowitz, S. D. & Regehr, W. G. Associative short-term synaptic plasticity mediated by endocannabinoids. Neuron 45, 419–431, doi:10.1016/j.neuron.2004.12.045 (2005).

56 Soler-Llavina, G. J. & Sabatini, B. L. Synapse-specific plasticity and compartmentalized signaling in cerebellar stellate cells. Nat Neurosci 9, 798–806, doi:10.1038/nn1698 (2006).

57 Lerner, T. N. & Kreitzer, A. C. RGS4 is required for dopaminergic control of striatal LTD and susceptibility to parkinsonian motor deficits. Neuron 73, 347–359, doi:10.1016/j.neuron.2011.11.015 (2012).

58 Gerdeman, G. L., Ronesi, J. & Lovinger, D. M. Postsynaptic endocannabinoid release is critical to long-term depression in the striatum. Nat Neurosci 5, 446–451, doi:10.1038/nn832 (2002).

59 Kreitzer, A. C. & Malenka, R. C. Endocannabinoid-mediated rescue of striatal LTD and motor deficits in Parkinson’s disease models. Nature 445, 643–647, doi:10.1038/nature05506 (2007).

60 Yasuda, H., Huang, Y. & Tsumoto, T. Regulation of excitability and plasticity by endocannabinoids and PKA in developing hippocampus. Proc Natl Acad Sci U S A 105, 3106–3111, doi:10.1073/pnas.0708349105 (2008).

61 Edwards, D. A., Zhang, L. & Alger, B. E. Metaplastic control of the endocannabinoid system at inhibitory synapses in hippocampus. Proc Natl Acad Sci U S A 105, 8142–8147, doi:10.1073/pnas.0803558105 (2008).

62 Li, B. Central amygdala cells for learning and expressing aversive emotional memories. Curr Opin Behav Sci 26, 40–45, doi:10.1016/j.cobeha.2018.09.012 (2019).

63 Katona, I. et al. Distribution of CB1 cannabinoid receptors in the amygdala and their role in the control of GABAergic transmission. J Neurosci 21, 9506–9518 (2001).

64 Morena, M., Patel, S., Bains, J. S. & Hill, M. N. Neurobiological Interactions Between Stress and the Endocannabinoid System. Neuropsychopharmacology 41, 80–102, doi:10.1038/npp.2015.166 (2016).

65 Gunduz-Cinar, O., Hill, M. N., McEwen, B. S. & Holmes, A. Amygdala FAAH and anandamide: mediating protection and recovery from stress. Trends Pharmacol Sci 34, 637–644, doi:10.1016/j.tips.2013.08.008 (2013).

66 Dana, H. et al. Sensitive red protein calcium indicators for imaging neural activity. Elife 5, doi:10.7554/eLife.12727 (2016).

67 Soltesz, I. et al. Weeding out bad waves: towards selective cannabinoid circuit control in epilepsy. Nat Rev Neurosci 16, 264–277, doi:10.1038/nrn3937 (2015).

68 Farrell, J. S. et al. In vivo assessment of mechanisms underlying the neurovascular basis of postictal amnesia. Sci Rep 10, 14992, doi:10.1038/s41598-020-71935-6 (2020).

69 Heinbockel, T. et al. Endocannabinoid signaling dynamics probed with optical tools. J Neurosci 25, 9449–9459, doi:10.1523/JNEUROSCI.2078-05.2005 (2005).

70 Ju, N. et al. Spatiotemporal functional organization of excitatory synaptic inputs onto macaque V1 neurons. Nat Commun 11, 697, doi:10.1038/s41467-020-14501-y (2020).

71 Sethuramanujam, S. et al. Rapid ‘multi-directed’ cholinergic transmission at central synapses. 2020.2004.2018.048330, doi:10.1101/2020.04.18.048330 %J bioRxiv (2020).

72 Kaifosh, P., Lovett-Barron, M., Turi, G. F., Reardon, T. R. & Losonczy, A. Septo-hippocampal GABAergic signaling across multiple modalities in awake mice. Nat Neurosci 16, 1182–1184, doi:10.1038/nn.3482 (2013).

73 Lovett-Barron, M. et al. Dendritic inhibition in the hippocampus supports fear learning. Science 343, 857–863, doi:10.1126/science.1247485 (2014).

74 Farrell, J. S. et al. Postictal behavioural impairments are due to a severe prolonged hypoperfusion/hypoxia event that is COX-2 dependent. Elife 5, doi:10.7554/eLife.19352 (2016).

75 Kaifosh, P., Zaremba, J. D., Danielson, N. B. & Losonczy, A. SIMA: Python software for analysis of dynamic fluorescence imaging data. Front Neuroinform 8, 80, doi:10.3389/fninf.2014.00080 (2014).

